# ATP-releasing SWELL1 channel in spinal microglia contributes to neuropathic pain

**DOI:** 10.1101/2023.01.08.523161

**Authors:** Jiachen Chu, Junhua Yang, Yuan Zhou, Jianan Chen, Kevin Hong Chen, Chi Zhang, Henry Yi Cheng, Nicholas Koylass, Jun O. Liu, Yun Guan, Zhaozhu Qiu

## Abstract

Following peripheral nerve injury, extracellular ATP-mediated purinergic signaling is crucial for spinal cord microglia activation and neuropathic pain. However, the mechanisms of ATP release remain poorly understood. Here, we show that volume-regulated anion channel (VRAC) is an ATP-releasing channel and is activated by inflammatory mediator sphingosine-1-phosphate (S1P) in microglia. Mice with microglia-specific deletion of Swell1 (also known as Lrrc8a), a VRAC essential subunit, had reduced peripheral nerve injury-induced increase in extracellular ATP in spinal cord. The mutant mice also exhibited decreased spinal microgliosis, dorsal horn neuronal hyperactivity, and both evoked and spontaneous neuropathic pain-like behaviors. We further performed high-throughput screens and identified an FDA-approved drug dicumarol as a novel and potent VRAC inhibitor. Intrathecal administration of dicumarol alleviated nerve injury-induced mechanical allodynia in mice. Our findings suggest that ATP-releasing VRAC in microglia is a key spinal cord determinant of neuropathic pain and a potential therapeutic target for this debilitating disease.

## INTRODUCTION

Neuropathic pain is a prevalent and debilitating chronic pain syndrome (*1*). It is often caused by injury and diseases that damage the nervous system. Spinal cord microglia, a type of resident immune cells in the central nervous system (CNS), play a critical role in the pathogenesis of neuropathic pain (*2, 3*). In response to peripheral nerve injury, spinal microglia are activated and undergo morphological changes, proliferation, and upregulation of signaling molecules, including many purinergic receptors (e.g. P2X4R, P2X7R, P2Y12R) (*4–6*). The purinergic system is one of the most important signaling pathways that regulate microglial activation (*7*). Upon stimulation of purinergic receptors, microglial cells release neuromodulators, which cause abnormal synaptic transmission and aberrant activity of spinal dorsal horn neurons (*8*). These pathological changes exaggerate spinal nociceptive transmission, and may also lead to the conversion of innocuous stimuli to painful signals. Blockage of purinergic receptors in the spinal cord rapidly reverses pain hypersensitivity (*5, 9*), indicating that neuropathic pain requires purinergic signaling triggered by extracellular ATP. However, the mechanisms of increased ATP release within the spinal cord following peripheral nerve injury remain elusive.

Activated by osmotic cell swelling, VRAC channel mediates the release of chloride (Cl^−^) and various organic osmolytes from the cells (*10*). This facilitates water efflux through osmosis and results in a regulatory volume decrease (*11*). ATP has also been detected from the supernatant of swollen cells in a VRAC-dependent manner using luciferin–luciferase assay (*12, 13*). However, it remains unclear whether ATP is released directly through VRAC itself or by indirect mechanisms. More importantly, the physiological relevance of VRAC-dependent ATP release remains to be determined since most mammalian cells in the body do not experience dramatic changes in extracellular osmolality. Interestingly, intrathecal injection of carbenoxolone, a nonspecific blocker of connexin/pannexin hemichannels and VRAC, was previously found to inhibit neuropathic pain in rodents, suggesting a potential pathological role for VRAC (*14–16*). Here, we show that VRAC is an ATP-permeable channel and mediates ATP release from microglia stimulated by S1P, an important inflammatory lipid mediator. Microglia-specific deletion of Swell1, the only obligatory VRAC subunit (*17, 18*), reduced nerve injury-induced increase of extracellular ATP content, attenuated microgliosis and neuronal hyperactivity in the spinal dorsal horn, and abolished neuropathic pain-like behaviors in mice. Using a high-throughput screening, we identified an FDA-approved drug dicumarol as a novel and potent VRAC inhibitor. Dicumarol administration in mice transiently alleviated neuropathic pain hypersensitivity. Collectively, our findings suggest the microglial VRAC channel as a key spinal determinant of neuropathic pain, which represents a potential novel therapeutic target for this debilitating chronic disease.

## RESULTS

### SWELL1-dependent VRAC is an ATP-releasing channel

To determine whether ATP directly permeates through the VRAC channel pore, we first performed experiments in HeLa cells expressing endogenous VRAC. We replaced intracellular Cl^−^ with equimolar ATP as the only permeant anion and perfused hypotonic solution to activate VRAC currents (*19*). Whole-cell patch-clamp recordings revealed substantial inward currents representing the efflux of negatively charged ATP from wild-type (WT) cells (**Fig. 1A**). Consistent with SWELL1 being the essential subunit of VRAC (*17, 18*), the inward currents mediated by ATP efflux were abolished in *SWELL1* knockout (KO) cells (**Fig. 1A**). This was rescued by overexpressing *SWELL1* cDNA, suggesting that SWELL1 is necessary for ATP permeability of VRAC (**Fig. 1A-1B**). The shifts in reversal potential indicate a significant ATP permeability of the channel (estimated P_ATP_/P_Cl_ at ∼0.06 assuming all intracellular ATP^4−^ as free tetravalent anions) (**Fig. 1C-1D**). We next performed the luciferin–luciferase assay and measured the bulk extracellular ATP released from intact HeLa cells after the treatment of hypotonic solution. In line with the electrophysiological data and findings from the previous knockdown experiments (*13*), cell swelling induced a dramatic increase in ATP efflux in WT cells, which was eliminated in *SWELL1* KO cells (**Fig. 1E**). This ATP release was also inhibited by a known VRAC blocker 4-(2-butyl-6, 7-dichloro-2-cyclopentyl-indan-1-on-5-yl) oxobutyric acid (DCPIB) in a dose-dependent manner (**Fig. 1E**).

**Fig. 1.**
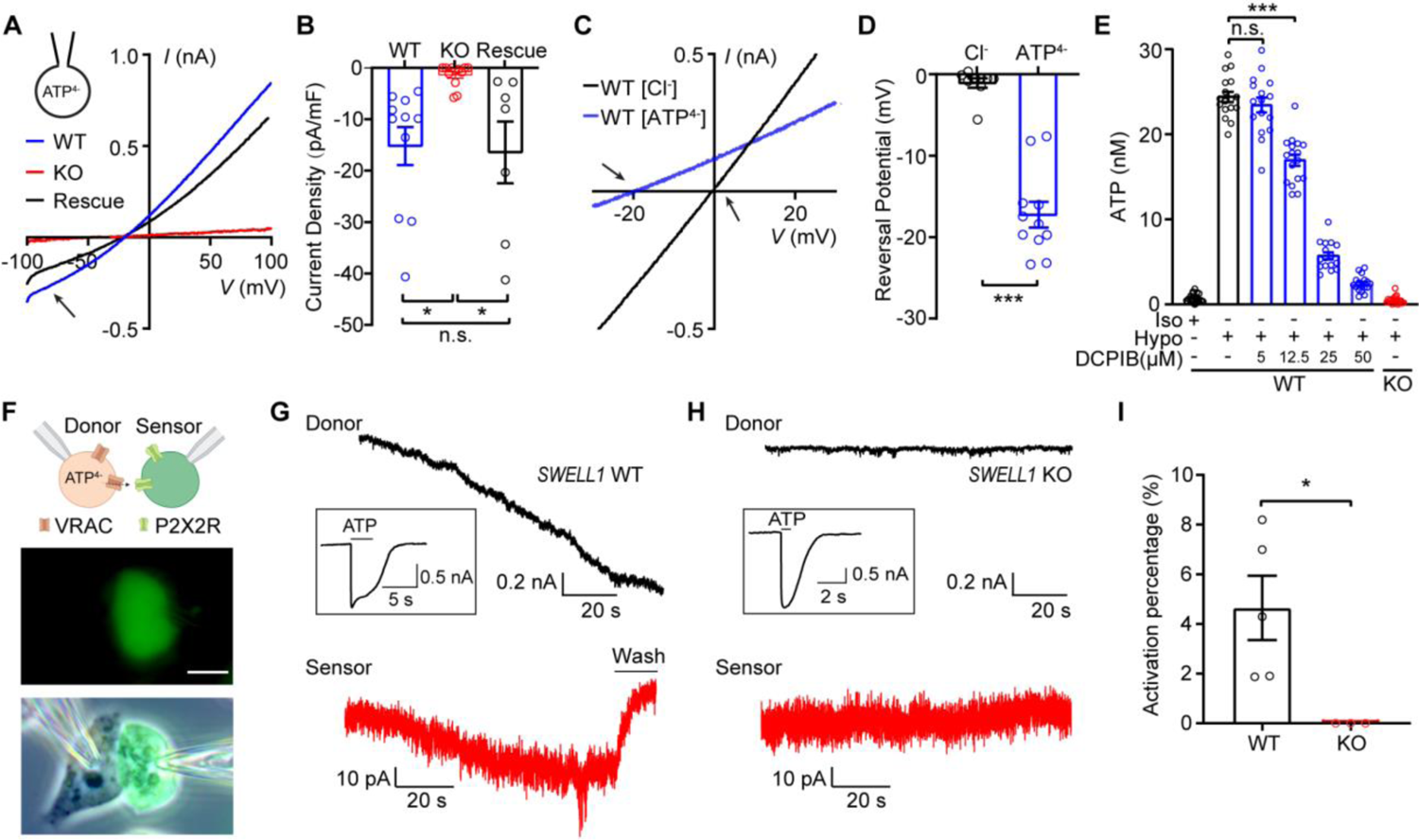
SWELL1-dependent VRAC is an ATP-releasing channel. (**A-B**) Hypotonicity (HYPO)-induced whole-cell currents (A) and their quantifications (B) at −100 mV with ATP-based pipette solution from WT and *SWELL1* KO HeLa cells as well as KO cells transfected with *SWELL1* cDNA (Rescue). Arrow indicates inward currents mediated by ATP^4-^ efflux. One-way ANOVA, Bonferroni post hoc test, n.s.: not significant, *p < 0.05. (**C-D**) HYPO-induced whole-cell currents (C) and their reversal potentials recorded by ramp protocol with Cl^−^-based or ATP^4−^-based pipette solution. Arrows indicate the reversal potentials. Student’s *t*-test, ***p < 0.001. (E) HYPO-induced ATP release in HeLa cells. n = 17 wells. One-way ANOVA, Bonferroni post hoc test, n.s.: not significant, ***p < 0.001. (F) Schematic illustration (top) and representative images (below) of sniffer-patch technique. Scale bar, 10 μm. (**G-H**) Representative current traces were recorded at −100 mV from (**G**) WT and (**H**) *SWELL1* KO HeLa cells, and their respective sensor cells (red). After each experiment, full receptor activations (inserted) in the sensor cells were recorded by direct ATP bath application. (**I**) Quantification of P2X2R activation. Student’s t-test, *p < 0.05 Bar graphs are reported as mean ± SEM.

To further test whether VRAC-mediated ATP release can be detected by surrounding cells, we employed the sniffer-patch method as a sensitive functional bioassay for extracellular ATP (*20*). As shown in **Fig. 1F**, we performed double whole-cell patch-clamp recordings in which HeLa cells, were the source cells, and the neighboring HEK293T cells transfected with P2X2R-YFP, a slowly-desensitizing ionotropic purinergic receptor, were the sensor cells. Hypertonic pipette solution (also causing osmotic cell swelling) with 10 mM ATP activated VRAC in WT source cells as indicated by the developing inward currents (**Fig. 1G**). Simultaneously, we observed inward currents in the neighboring sensor cells, suggesting the activation of P2X2R (**Fig. 1G**). To quantify the amount of ATP release, we normalized P2X2R currents to the maximal receptor activation by direct bath application of 10 mM ATP after each double recording experiment (**Fig. 1G-1I**). This revealed significant VRAC-dependent ATP efflux (∼5% of full P2X2R activation) detected by the adjacent sensor cells (**Fig. 1I**). As expected, VRAC currents were abolished in *SWELL1* KO source cells (**Fig. 1H**), and ATP release from KO cells was also absent (**Fig. 1H-1I**). Together, these data provide direct evidence that SWELL1-dependent VRAC channel conducts and releases ATP.

### Swell1 is essential for VRAC activity in microglia and mediates ATP release

To determine whether Swell1 is required for VRAC activity in microglia, we first performed western blot and observed robust Swell1 protein expression in mouse BV2 microglial cells (**Fig. 2A**). Perfusion of hypotonic solution elicited prominent Cl^−^ currents in microglia with characteristic features of VRAC, including outward rectification and DCPIB sensitivity (**Fig. 2B-2D**). In addition to cell swelling, VRAC can also be activated by other physiologically relevant stimuli, including S1P (*21*), an inflammatory mediator implicated in neuropathic pain (*22*). Indeed, S1P in normal isotonic buffer induced DCPIB-sensitive outwardly-rectifying VRAC currents in microglia (**Fig. 2E-2G**) (*23*). Further pharmacological studies suggest that S1P-induced VRAC activity in microglia is dependent on S1P receptors (S1PR1 and S1PR2) and the production of reactive oxygen species (ROS), but not Ca^2+^ (**Figure S1**). Consistently, S1P also induced robust ATP release from microglia (**Fig. 2H**). To test whether S1P-induced ATP release is dependent on VRAC, we generated *Swell1* KO BV2 microglia cells using CRISPR-Cas9 technology (**Fig. 2A**). The loss of Swell1 protein abolished both VRAC currents and ATP efflux upon S1P treatment (**Fig. 2B-2H**), suggesting that Swell1 is an essential subunit of VRAC in BV2 microglia and may mediate ATP release during inflammation.

**Fig. 2.**
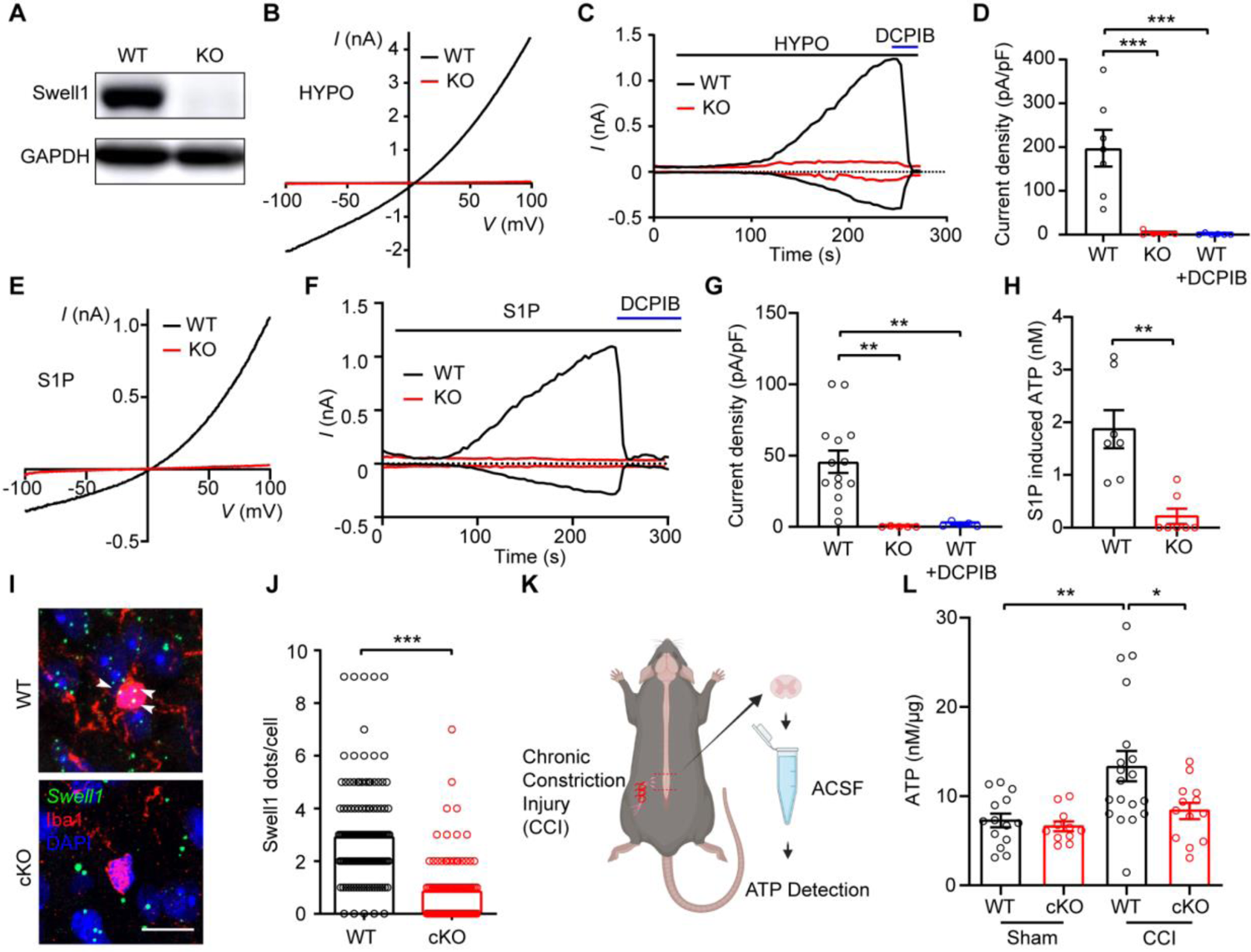
Microglial Swell1 is essential for hypotonicity- and S1P-induced VRAC activation and mediates ATP release. (A) Representative western blots showing the loss of Swell1 protein from *Swell1* KO BV2 microglia. (B) Representative HYPO-induced whole-cell currents recorded by ramp protocol for WT and *Swell1* KO BV2 microglia. (**C-D**) Time course at ±100 mV (C) and quantification at +100 mV (D) of HYPO-induced whole-cell currents for WT and *Swell1* KO BV2 microglia. DCPIB was applied at 50 μM. One-way ANOVA, Bonferroni post hoc test, n.s.: not significant, ***p < 0.001. (**E**) Representative whole-cell currents recorded by ramp protocols induced by 10 nM S1P for BV2 microglia. (**F-G**) Time course at ±100 mV (F) and quantification at +100 mV (G) of S1P-induced whole-cell currents BV2 microglia. One-way ANOVA, Bonferroni post hoc test, n.s.: not significant, **p < 0.01. (**H**) S1P-induced ATP release from BV2 microglia. Student’s *t*-test, *p < 0.01. (**I-J**) Representative images and quantification of *Swell1* RNAscope *in situ* hybridization in spinal cord dorsal horns. Note that mRNA dots (white arrows) were counted only when they colocalize with Iba1-positive microglial soma. Due to the small soma size, *Swell1* mRNA expression is probably underestimated. Scale bar, 10 μm. n = 140 microglia cells from 4 mice for WT; n = 108 microglia cells from 4 mice for *Swell1* cKO. Student’s *t*-test, ***p < 0.001. (K) Schematic diagram of ATP detection following CCI. (L) ATP levels in ACSF from lumbar spinal cord slices. Two-way ANOVA, Bonferroni post hoc test, *p < 0.05, **p < 0.01. Bar graphs are reported as mean ± SEM.

Extracellular ATP level is known to increase in the spinal cord after peripheral nerve injury (*24*). To examine whether microglia VRAC channel contributes to this ATP enhancement, we generated microglia-specific *Swell1* conditional knockout mice (cKO) by breeding *Swell1^F/F^* mice with the *Cx3cr1-Cre* line, a widely used genetic tool for investigating microglia function (*25*). Due to the lack of specific antibody for immunostaining, we performed RNAscope *in situ* hybridization and validated the decrease of *Swell1* mRNA expression in spinal microglia in the cKO mice (**Fig. 2I-2J**). We then carried out the well-established chronic constriction injury (CCI) model of neuropathic pain by tying three loose monofilament ligatures around the unilateral sciatic nerve (**Fig. 2K**) (*26*). As expected, the concentration of extracellular ATP in the artificial cerebral spinal fluid (ACSF) incubated with L3-5 spinal cord slices from *Swell1^F/F^* WT control mice was significantly higher in the CCI group than the sham surgery group (**Fig. 2L**). Interestingly, the CCI-induced increase in extracellular ATP was dampened in slices prepared from *Swell1* cKO mice (**Fig. 2L**). These results suggest that microglia contribute to the increased ATP release after peripheral nerve injury via a Swell1 channel-dependent mechanism. The reduction in extracellular ATP levels in the mutant mice could be due to the decreased ATP release through microglial Swell1 channel directly, or the reduced microglial activation indirectly (see data below), or more likely both.

### Reduction of CCI-induced neuropathic pain-like behaviors in microglia-specific *Swell1* cKO mice

To test if microglial Swell1 expression is regulated by peripheral nerve injury, we performed RNAscope *in situ* hybridization and observed that *Swell1* mRNA level in microglia was increased in the spinal cord dorsal horn after sciatic CCI (**Fig. S2**). This indicates that microglial Swell1 expression is upregulated in response to nerve injury. To determine whether Swell1 is involved in the pathogenesis of neuropathic pain, we first evaluated tactile allodynia in microglia-specific *Swell1* cKO male mice after CCI by investigators blind to the mouse genotype. The ipsilateral paw withdrawal frequencies evoked by innocuous mechanical stimuli with two von Frey filaments in *Swell1^F/F^* WT male mice were significantly increased after CCI (**Fig. 3A-3C**). Such increases were not observed in *Swell1* cKO male mice (**Fig. 3A-3C**), suggesting that microglial Swell1 is critical for the development of neuropathic mechanical hypersensitivity in males. We then accessed the extent of CCI-induced thermal hyperalgesia using the Hargreaves test. The ipsilateral paw withdrawal latencies of WT male mice were significantly decreased after CCI, suggesting the development of heat hypersensitivity (**Fig. 3D**). Again, such change was absent in cKO mutants (**Fig. 3D**), indicating that microglia Swell1 also contributes to nerve injury-induced heat hypersensitivity in male mice. Importantly, before the CCI surgery, *Swell1* cKO male mice and their control littermates exhibited similar responses to mechanical and heat stimuli (**Fig. 3B-3D**), suggesting that loss of microglia Swell1 does not cause defects in nociceptive pain sensitivity under normal conditions.

**Fig. 3.**
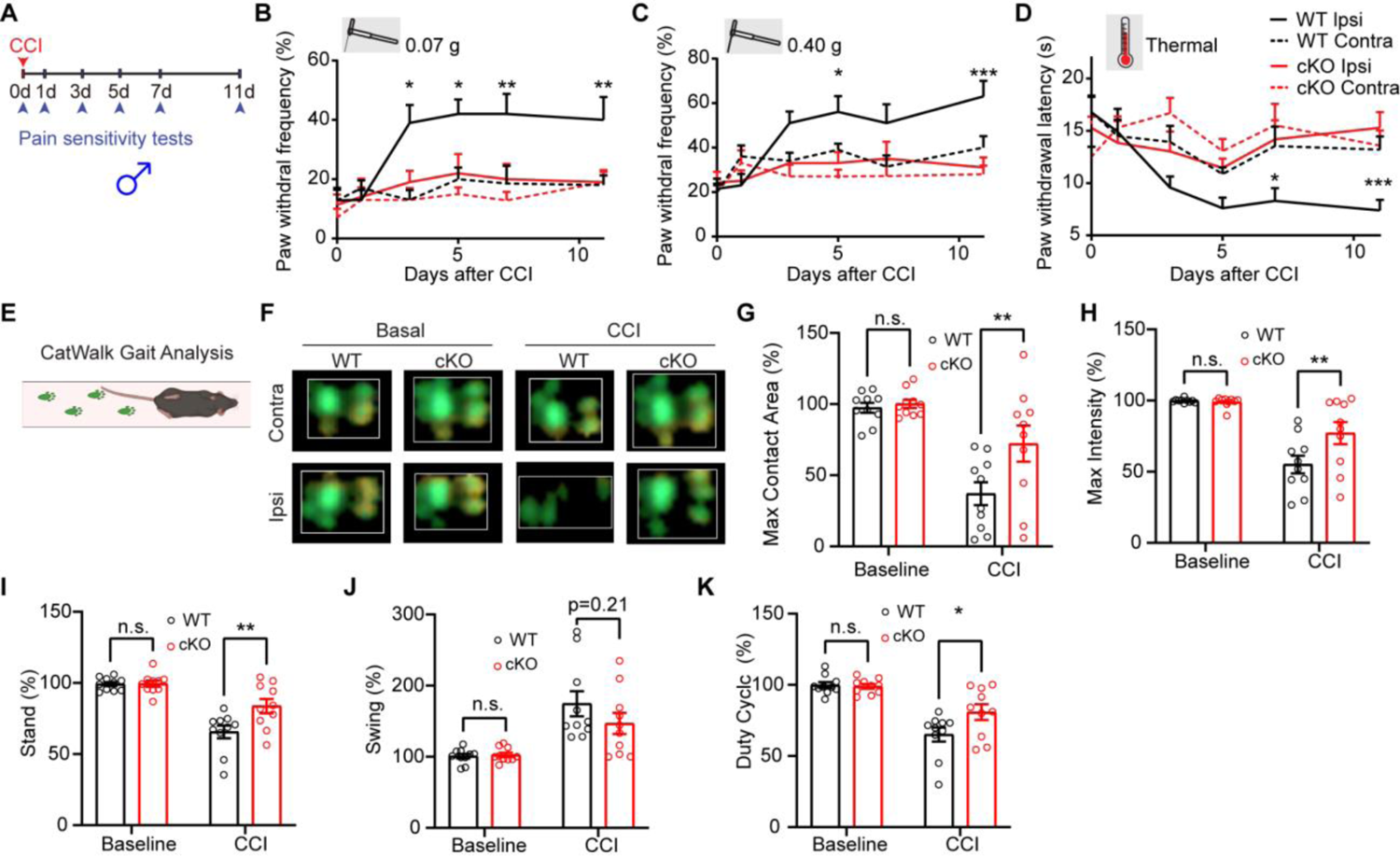
Genetic deletion of microglial Swell1 alleviates CCI-induced neuropathic pain-like behaviors in male mice. (**A**) Experimental design of measuring CCI-induced neuropathic pain-like behaviors. Baseline sensitivity was measured at day 0 before CCI. (**B-D**) Baseline and CCI-induced paw withdrawal frequency detected by the von Frey filaments (B: 0.07 g; C: 0.40 g) and paw withdrawal latency detected by Hargreaves test (D) in WT (n = 10) and *Swell1* cKO (n = 10) mice from ipsilateral and contralateral sides. Statistics between WT ipsilateral side and *Swell1* cKO ipsilateral side were labeled. Two-way ANOVA, Bonferroni post hoc test, *p < 0.05, **p < 0.01, ***p < 0.001. (E) Schematic of the CatWalk gait analysis system. (F) Representative combined paw print images in CatWalk assay. **(G-K)** Quantification of the right hind paw (ipsilateral to the side of nerve injury) maximum contact area (G), intensity (H), stand (I), swing (J), duty cycle (K) normalized to left hind paw (contralateral to the side of nerve injury) at baseline and 7 days after CCI in WT mice (n = 10) and *Swell1* cKO mice (n = 10). Two-way ANOVA, Bonferroni post hoc test, n.s.: not significant, *p < 0.05, **p < 0.01. Bar graphs are reported as mean ± SEM.

In addition to measuring reflex pain behaviors evoked by mechanical and heat stimuli, we adopted an automated CatWalk gait analysis to examine changes in pain-related gait parameters after CCI. This quantitative gait analysis represents an objective way to examine changes in various walking parameters due to pain and motor impairment and has been successfully used to assess locomotor deficits and pain syndromes in animal models (*27, 28*). CatWalk measures the pressure map produced by the paws using an optics-based pressure sensor that detects distortions on a glass platform (**Fig. 3E-3F**). In freely moving, non-restrained mice, we measured several parameters, including maximum contact area and intensity of a paw that comes into contact with the glass during walking, stand (the duration of contact), swing (the duration of no contact) of a paw with the glass plate, and duty cycle: (stand / (stand + swing) × 100%. To exclude the influence of other confounding factors such as body weight and paw size, we normalized the paw parameters from the ipsilateral hind paw to those from the contralateral side and expressed data as percentages to reflect the gait abnormalities. Previous studies show that animals suffering from pain after nerve injury would reduce weight support upon the affect hindpaw/hindlimb during walking, due to ongoing pain and movement-induced pain (*27, 28*). The changes in gait parameters correlated with other pain symptoms, such as mechanical allodynia. Indeed, maximum contact area and intensity, stand and duty cycle of the ipsilateral paw were decreased, and swing was increased in WT male mice one week after CCI (**Fig. 3G-3K**). Interestingly, these changes were significantly reduced in *Swell1* cKO males except swing, the difference of which trends toward statistical significance (**Fig. 3G-3K**). Since cKO mice did not exhibit any general defects in locomotor function in the CatWalk assay and open field test (**Fig. S3**), these data suggest that CCI-induced gait abnormalities were reduced in the absence of microglial Swell1 in males. Together, our animal behavioral studies demonstrate that Swell1 channel in microglia plays a key role in the pathogenesis of neuropathic pain-like behaviors in male mice.

Sexual dimorphism of microglia and its role in neuropathic pain has been well-documented (*2, 3*). To examine if there is such a sexual difference for microglial Swell1, we performed the same battery of behavioral tests in female mice following peripheral nerve injury (**Fig. S4**). Compared to WT female controls, *Swell1* cKO female mice exhibited significant alleviation from CCI-induced pain hypersensitivity in some assays (0.07 g von Frey test; maximal contact area and swing in CatWalk). However, the extent of the protection from pain-like behaviors in other assays was less obvious compared to that in the mutant male mice (**Fig. S4**). These data suggest an interesting sexual dimorphism of microglial Swell1 in regulating pain-like behaviors with a more prominent role in male mice than female mice. This sexual difference is consistent with some recent literature regarding a uniquely important role of microglia in pain in male but not female mice (*29*). The underlying mechanisms warrant further investigation.

### Reduction of CCI-induced spinal microgliosis and dorsal horn neuronal hyperactivity in microglia-specific *Swell1* cKO mice

In response to peripheral nerve injury, microglia in the spinal cord undergo robust proliferation and morphological changes (*2, 3*), which are critical for the development of neuropathic pain. To examine CCI-induced microglia activation, we performed immunostaining with microglia marker Iba1 in L3-5 spinal cord sections. In the spinal dorsal horn of sham surgery group, microglia exhibited the typical ramified morphology and a normal density in both WT and *Swell1* cKO mice (**Fig. 4A**), suggesting that loss of Swell1 does not cause abnormal microglia activation under physiological conditions. As expected, we observed obvious spinal microgliosis in WT control mice following CCI, based on a marked increase in Iba-1 immunoreactivity (**Fig. 4A-4C**). However, these changes induced by CCI were reduced in cKO mice (**Fig. 4A-4C**), indicating that Swell1 is required for CCI-induced cellular alterations of spinal microglia. Activated spinal microglia are known to secrete various signaling molecules including interleukin-1β (IL-1β), a key proinflammatory cytokine, which drives dorsal horn neuron hypersensitivity and neuropathic pain (*30*). Consistent with the reduced microglial activation, spinal cord IL-1β concentration was also lower in *Swell1* cKO mice following CCI compared to WT controls (**Fig. 4D**). Together, these results demonstrate that microglial Swell1 is an important regulator for microglia activation and proinflammatory changes occurred in the spinal cord following nerve injury.

**Fig. 4.**
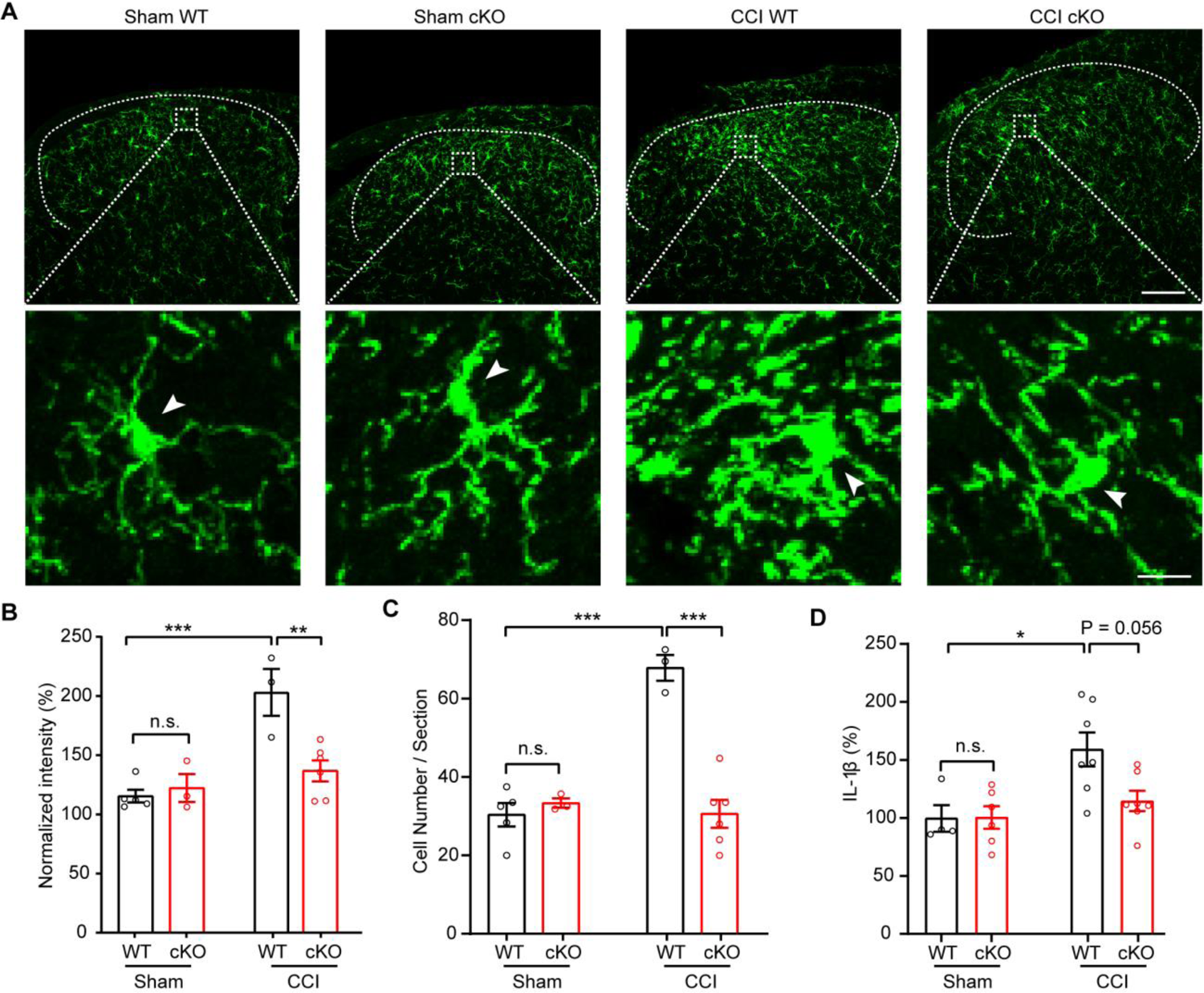
CCI-induced microgliosis is reduced in microglia-specific *Swell1* cKO mice. (A) Confocal immunofluorescence microscopy of representative spinal cord sections showing microglia (Iba1 staining) in Sham and CCI-treated WT and *Swell1* cKO mice. Upper scale bar, 100 μm; lower scale bar, 10 μm. Arrows indicate the microglia cell soma. (B) Quantification of Iba1 staining intensity in spinal cord L3-L5 dorsal horn. Ipsilateral Iba1 intensity was normalized to the contralateral side of each section (data from 3 sections per mice were averaged). n = 5 mice for Sham WT; n = 3 mice for Sham *Swell1* cKO; n = 3 mice for CCI WT; n = 6 mice for CCI *Swell1* cKO. Two-way ANOVA, Bonferroni post hoc test, n.s.: not significant, **p < 0.01, ***p < 0.001. (C) Quantification of microglial cell number per spinal cord L3-L5 dorsal horn section (data from 3 sections per mice were averaged). n = 5 mice for Sham WT; n = 3 mice for Sham *Swell1* cKO; n = 3 mice for CCI WT; n = 6 mice for CCI *Swell1* cKO. Two-way ANOVA, Bonferroni post hoc test, n.s.: not significant, ***p < 0.001. (D) Quantification of IL-1β levels in L3-L5 spinal cord total lysate measured by ELISA. n = 4 mice for Sham WT; n = 6 mice for Sham *Swell1* cKO; n = 7 mice for CCI WT; n = 7 mice for CCI *Swell1* cKO. Two-way ANOVA, Bonferroni post hoc test, n.s.: not significant, *p < 0.05. Bar graphs are reported as mean ± SEM.

VRAC-mediated Cl^−^ efflux has been previously proposed as an important step in NLRP3 (NOD-, LRR- and pyrin domain-containing protein 3) inflammasome activation, leading to the secretion of IL-1β from macrophages (*31*). To test whether Swell1 is directly involved in NLRP3 activation and IL-1β release, we cultured bone marrow-derived macrophages (BMDM) from *Swell1* cKO mice and validated the loss of VRAC currents by whole-cell patch-clamp recordings (**Fig. S5A-S5B**). Upon stimulation with various damage-associated molecular patterns (DAMP) including ATP and nigericin, *Swell1* KO BMDMs released similar levels of IL-1β as WT cells (**Fig. S5C-S5D**). A recent study reported that Swell1 is essential for hypotonicity-induced NLRP3 inflammasome activation (*32*). However, we did not observe a defect in hypotonicity-induced IL-1β release from *Swell1* KO BMDMs. These data suggest that Swell1 is dispensable for NLRP3 inflammasome activation and that VRAC regulates IL-1β levels in the spinal cord likely through an indirect mechanism involving the modulation of microglia activation.

Microglia-neuron interactions play a crucial role in regulating neuropathic pain characterized by increased activity of dorsal horn neurons (*2, 3*). To test whether microglial Swell1 regulates this process, we first performed immunostaining of c-Fos, an immediate early gene product and a classical molecular marker of neuronal activation. There was a marked increase in the number of c-Fos positive (+) neurons within the spinal dorsal horn of WT mice after CCI (**Fig. 5A-5B**), which may be partly due to increases of spontaneous neuronal activity and neuronal activation during exploration in awake state after nerve injury. Comparatively, the number of c-Fos+ dorsal horn neurons was significantly less in *Swell1* cKO mice following CCI (**Fig. 5A-5B**). To assess synaptic transmission, we next performed whole-cell patch-clamp recordings on acute spinal cord slices in a genotype-blinded manner and measured spontaneous excitatory postsynaptic currents (sEPSCs) in lamina II dorsal horn neurons which are crucial in processing nociceptive information (*2, 3*). Although lamina II neurons are heterogenous, previous studies showed that nerve injury generally increases their synaptic transmission and excitability (*33*). Indeed, we observed that the amplitude and frequency of sEPSCs in lamina II neurons were markedly increased in WT mice after CCI, as compared to those after sham surgery (**Fig. 5C-5E**). Interestingly, CCI-induced increases in sEPSC amplitude and frequency were both dampened in *Swell1* cKO mice (**Fig. 5C-5E**). Collectively, these findings suggest that microglial Swell1 channel contributes to CCI-induced dorsal horn neuronal hyperactivity and the enhancement of excitatory synaptic transmission.

**Fig. 5.**
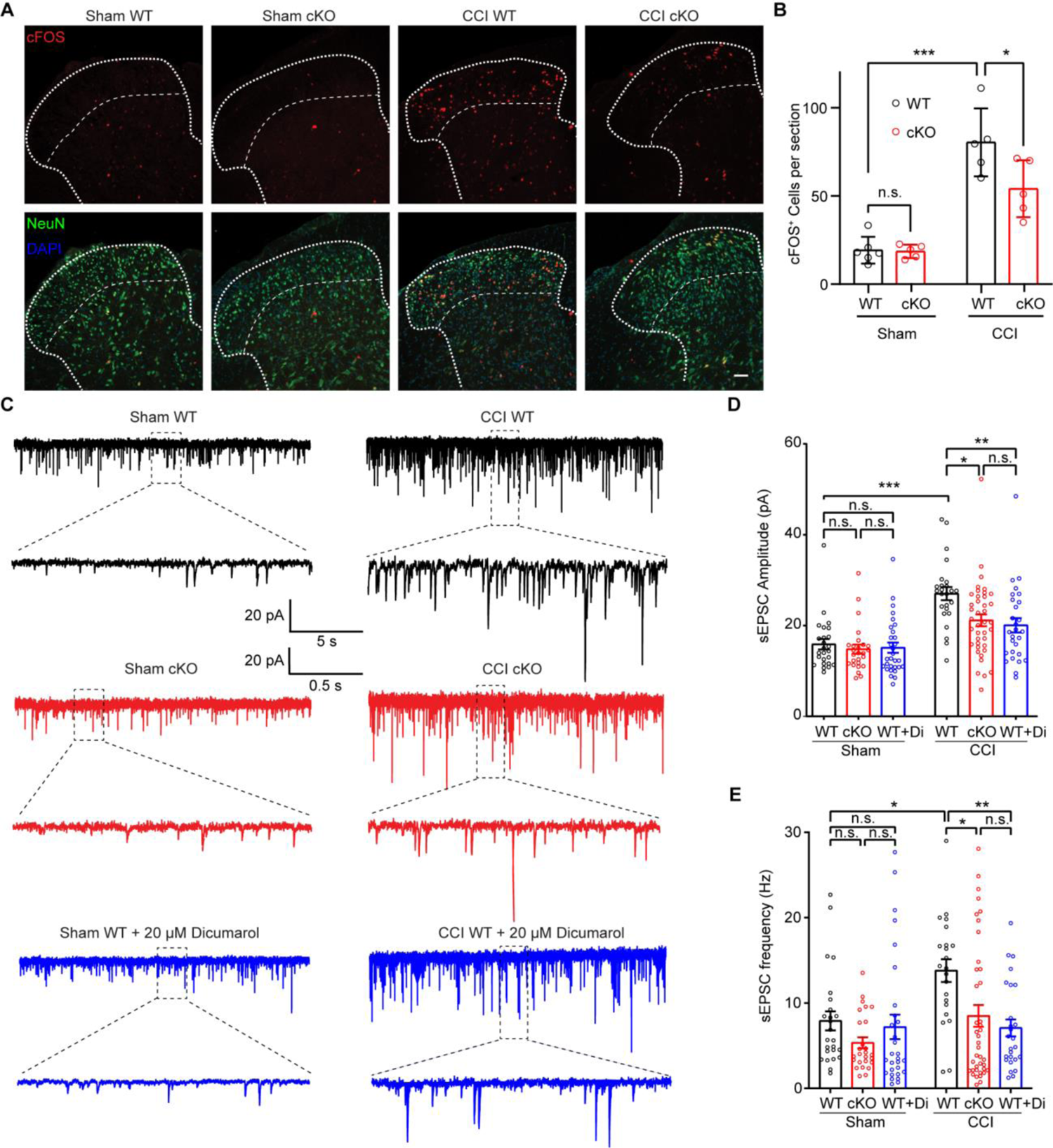
Swell1 channel contributes to CCI-induced dorsal horn neuronal hyperactivity and enhanced excitatory synaptic transmission. (**A**) Representative images and (**B**) Quantifications of c-Fos immunostaining in superficial laminae of spinal cord sections from WT and *Swell1* cKO mice with Sham or CCI treatment. n = 6 mice for Sham WT; n = 5 mice for Sham *Swell1* cKO; n = 5 mice for CCI WT; n = 5 mice for CCI *Swell1* cKO (data from 3 sections per mice were averaged). Two-way ANOVA, n.s.: not significant, Bonferroni post hoc test, *p < 0.05, ***p < 0.001. Scale bar, 50 μm. (**C-E**) Representative traces (C), and quantifications of amplitude (D) and frequency (E) of sEPSCs recorded from lamina II dorsal horn neurons of acute spinal cord slices. n = 25 cells from 3 Sham WT mice, n = 26 cells from 3 Sham cKO mice, n = 30 cells from 3 Sham WT mice with 20 μM dicumarol incubation, n = 25 cells from 3 CCI WT mice, n = 40 cells from CCI cKO mice, n = 28 cells from 3 CCI WT mice with 20 μM dicumarol incubation. Two-way ANOVA, n.s.: not significant, Bonferroni post hoc test, *p < 0.05, **p < 0.01, ***p < 0.001. Bar graphs are reported as mean ± SEM.

### An FDA-approved drug library screen identifies dicumarol as a potent VRAC inhibitor

To further support the genetic studies, we sought to investigate the functional role of VRAC in neuropathic pain with pharmacological inhibitors. The currently available VRAC small-molecule blockers often lack potency and specificity (*34*). To search for novel VRAC inhibitors, we employed a high-throughput yellow fluorescent protein (YFP) quenching assay. This assay was developed for our unbiased RNAi screens, which led to the successful identification of SWELL1 as an essential component of VRAC (*17*) and PAC as the proton-activated Cl^−^ channel (*35*). Treatment of hypotonic solution to HEK293T cells stably expressing an iodide-sensitive YFP activates the endogenous VRAC channel (**Fig. 6A**). A subsequent addition of iodide (I^−^) triggers a rapid fluorescence decrease due to I^−^ influx through VRAC (**Fig. 6B**). To identify small molecules with translational potential, we performed the YFP quenching screen with a library containing ∼3000 existing drugs (**Fig. 6C**) (*36*). We followed up the potential hits (>2 standard deviations above the mean) with a secondary screen and electrophysiological recordings (**Fig. S6A-S6B**). This resulted in the identification of two new VRAC blockers: dicumarol, a competitive inhibitor of vitamin K epoxide reductase, and zafirlukast, a cysteinyl leukotriene receptor 1 (CysLT1R) antagonist. Application of dicumarol and zafirlukast led to rapid full inhibition of hypotonicity-induced VRAC currents in a voltage-independent and reversible manner (**Fig. 6D-6E** and **Fig. S6C**). The inhibitory effects of dicumarol and zafirlukast were dose-dependent with half-maximal inhibitory concentrations (IC_50_) of 3.8 and 4.9 µM, respectively (**Fig. 6F** and **Fig. S6D**). Notably, zafirlukast has also recently been found to inhibit VRAC from another drug screen (*37*). In comparison with zafirlukast, dicumarol is slightly more potent. Importantly, however, the maximum serum concentration (C_max_) for dicumarol (∼59 µM (*38*); National Center for Advancing Translational Sciences Inxight Drugs) far exceeds its IC_50_, suggesting significant inhibition of VRAC channel activity can be achieved using existing drug dosing regimen in humans. In contrast, the C_max_ for zafirlukast is at sub-µM concentrations (*39*), far below its IC_50_ value. We thus decided to focus on dicumarol for further characterization.

**Fig. 6.**
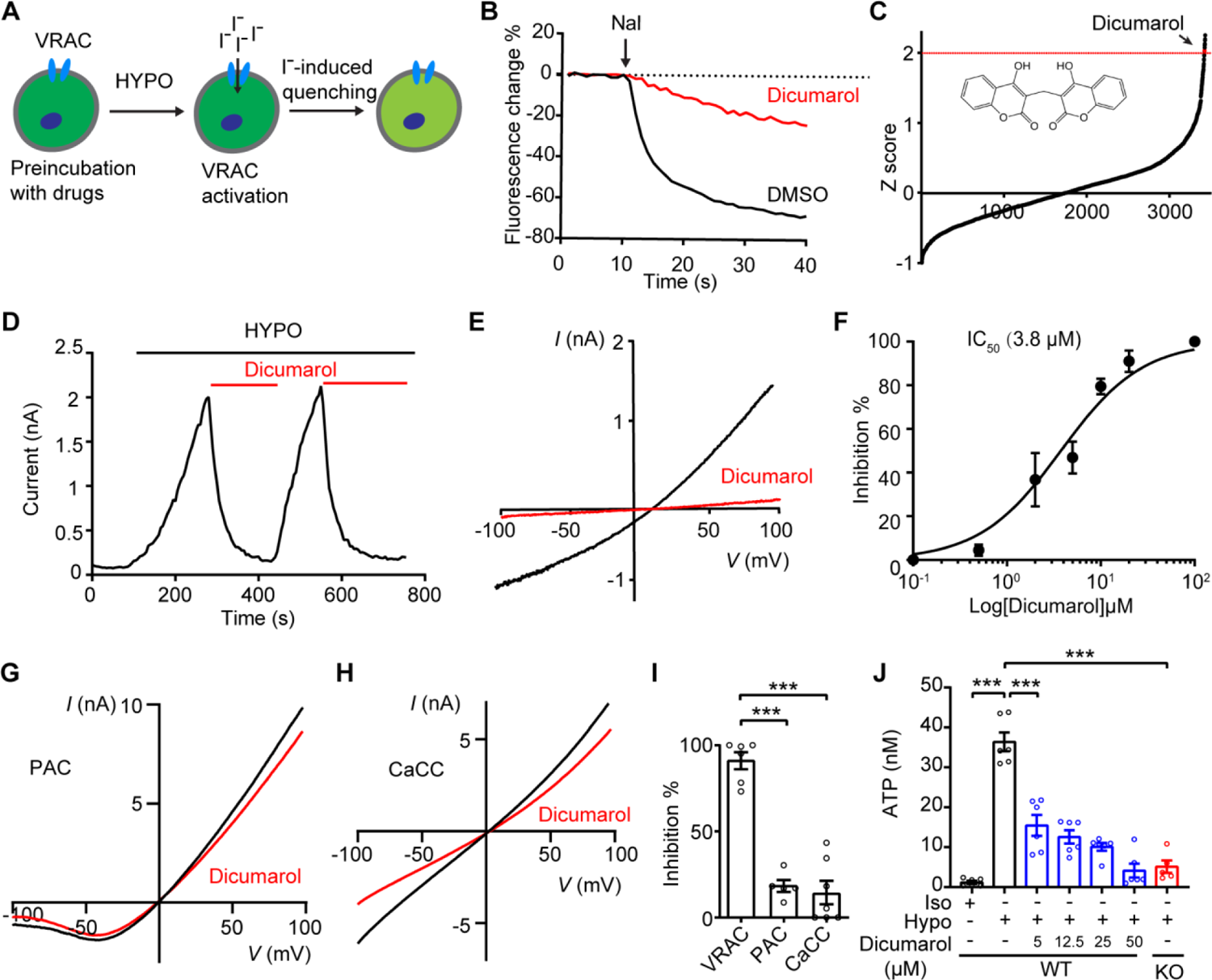
An FDA-approved drug dicumarol is a potent and specific VRAC inhibitor. (A) Schematic illustration of YFP quenching assay for VRAC inhibitor screening. (B) Representative traces of YFP quenching assay in HEK293-YFP cells treated with 20 µM dicumarol or DMSO control. (C) Z-score (the number of SD from the mean) of FDA-approved drugs plotted according to the rank order. Dicumarol is highlighted in red. (**D** and **E**) Hypotonicity-induced currents at +100 mV (D) and with voltage ramp (E) were inhibited by 10 µM dicumarol in HEK293T cells. HYPO: 200 mOsm/kg. (F) Dose-inhibition curve of dicumarol on hypotonicity-induced VRAC currents in HEK293T cells (n = 5-7 cells for each concentration). (G) Representative acid (pH 5.0)-induced whole-cell currents in HEK293T cells overexpressing human PAC with or without 20 µM dicumarol treatment. (H) Representative Ca^2+^(1 µM)-activated whole-cell currents in HEK293T cells overexpressing mouse Tmem16a with or without 20 µM dicumarol treatment. (I) Quantification of dicumarol-mediated inhibitions on three chloride channels. n = 5-7 cells for each group. One-way ANOVA, Bonferroni post hoc test, ***p < 0.001. (J) Hypotonicity-induced ATP release from HeLa cells were inhibited by dicumarol. n = 5-6 replicates for each group. One-way ANOVA, Bonferroni post hoc test, ***p < 0.001. Bar graphs are reported as mean ± SEM.

Originally isolated from molding sweet-clover hay, dicumarol is the prototype of the hydroxycoumarin anticoagulant drug class that depletes vitamin K in the blood (*40*). To determine whether the inhibitory effect of dicumarol on VRAC is dependent on its anticoagulant activity, we tested warfarin and 4-hydroxycoumarin, two anticoagulants of the same family (*40*), with patch-clamp recordings. Both of them failed to inhibit hypotonicity-induced VRAC currents, suggesting that dicumarol blocks the VRAC channel independent of its effect on vitamin K epoxide reductase (**Fig. S6E-S6F**). Dicumarol is a dimer of 4-hydroxycoumarin linked by a methylene bridge. These results also indicate that dimeric structure of coumarin is critical for blocking the VRAC channel.

The existing VRAC blockers lack specificity and often cross-inhibit other families of Cl^−^ channels (*34*). To test if dicumarol is selective for VRAC, we expressed PAC channel and Ca^2+^-activated Cl^−^ channel (CaCC/TMEM16A) in HEK293T cells and observed no obvious inhibitory effect of dicumarol on these two channels (**Fig. 6G-6I**). These data indicate that dicumarol is a relatively specific VRAC inhibitor. Consistent with its strong inhibitory effect on VRAC currents, dicumarol also blocked hypotonicity-induced ATP release in a dose-dependent manner (**Fig. 6J**). Notably, while 5 µM DCPIB was not effective (**Fig. 1E**), dicumarol at this concentration already suppressed ATP release (**Fig. 6J**). Together, our results demonstrate that dicumarol, an existing FDA-approved drug, is a potent and selective VRAC inhibitor.

### Blocking the VRAC channel with dicumarol alleviates CCI-induced mechanical hypersensitivity

To examine whether dicumarol can inhibit VRAC activity in microglia, we activated VRAC currents in BV2 microglia with hypotonic solution and then treated them with dicumarol. Application of dicumarol in microglia inhibited hypotonicity-induced VRAC currents in a dose-dependent manner with a similar IC_50_ (4.1 µM) as that in HEK293T cells (**Fig. 7A-7B**). Additionally, dicumarol treatment effectively blocked S1P-induced VRAC currents and ATP release from microglia (**Fig. 7C-7E**). These data indicate that dicumarol is also a potent inhibitor for VRAC activated by inflammatory stimuli in microglia.

**Fig. 7.**
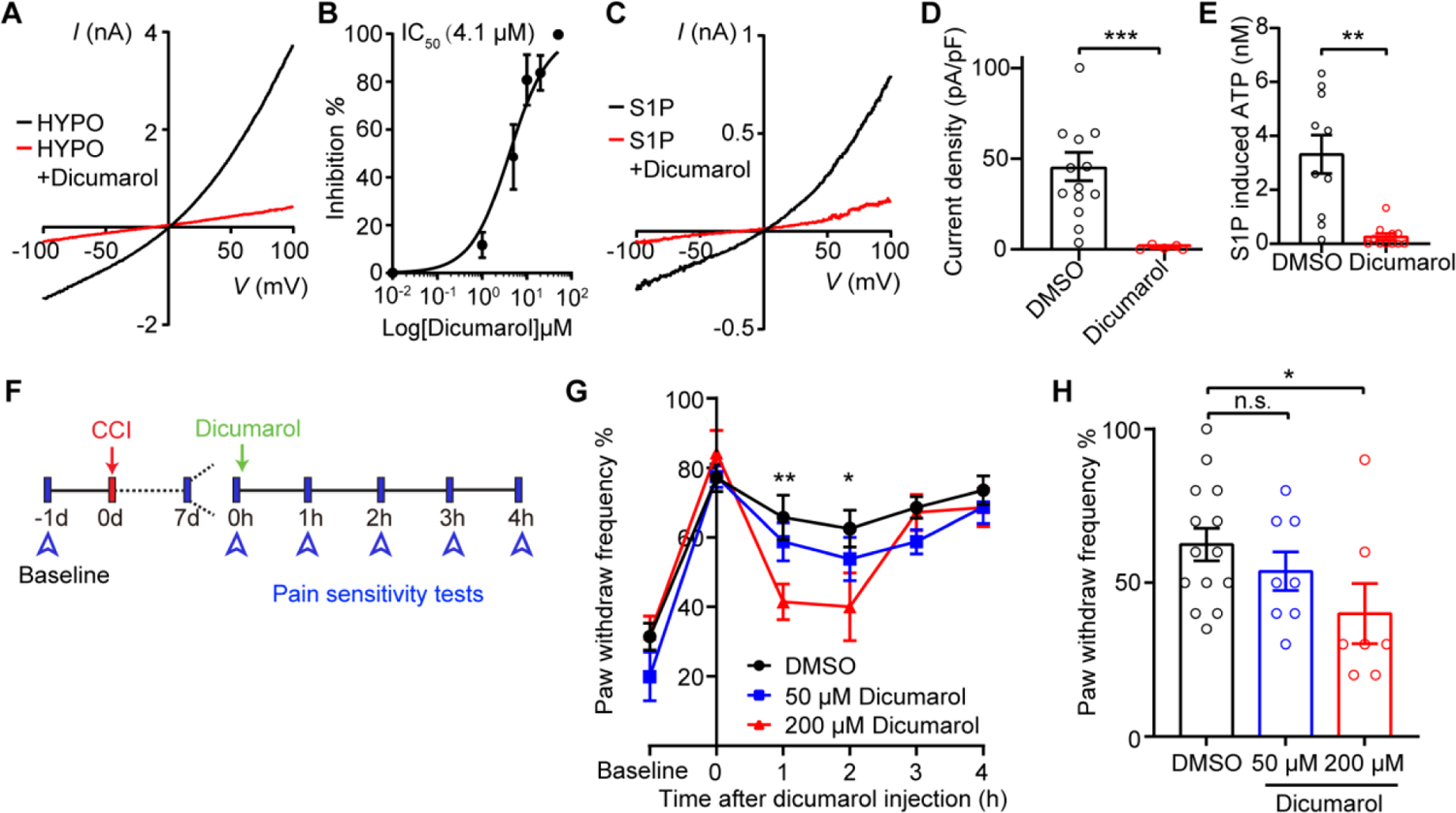
Dicumarol reduces CCI-induced mechanical pain hypersensitivity. (A) Representative hypotonicity (HYPO)-induced VRAC currents in BV2 microglia with or without 20 µM dicumarol treatment. (B) Dose-inhibition curve of dicumarol on hypotonicity-induced VRAC currents in BV2 microglia (n = 5-7 cells for each concentration). (C) Representative S1P (10 nM)-induced VRAC currents in BV2 microglia with or without 20 µM dicumarol treatment. (D) Quantification of S1P-induced VRAC current densities at +100 mV treated with DMSO (n = 14) or dicumarol (n = 5). Student’s *t* tests, ***p < 0.001. (E) 10 nM S1P-induced ATP release from BV2 microglia treated with DMSO or 10 µM dicumarol. n = 10 replicates from 4 independent experiments for DMSO, and n=11 replicates from 4 independent experiments for dicumarol. Student’s *t* tests, **p < 0.01. (F) Experimental design of dicumarol treatment in CCI-induced neuropathic pain mouse model. (**G** and **H**) Paw withdrawal frequency was detected by the von Frey filament (0.40 g) in CCI-treated WT mice before and after intrathecal injection of 5 µL DMSO or dicumarol. Statistics between DMSO and 200 µM dicumarol were labeled. Paw withdrawal frequency 2 hours after injection are plotted in (H). n = 14 mice for DMSO, n = 8 mice for 50 µM dicumarol, n = 7 mice for 200 µM dicumarol. Two-way ANOVA, Bonferroni post hoc test, n.s.: not significant, *p < 0.05, **p < 0.01. Bar graphs are reported as mean ± SEM.

To further test whether inhibiting VRAC with dicumarol suppresses nerve injury-induced neuronal hyperactivity, we incubated spinal cord slices with dicumarol before whole-cell patch-clamp recordings. Dicumarol treatment markedly inhibited CCI-induced increases in sEPSC amplitude and frequency in lamina II neurons of WT mice (**Fig. 5C-5E**). Based on these results, we further performed a proof-of-concept study to test if dicumarol attenuates neuropathic pain-like behaviors. Intrathecal administration of 5 µL dicumarol at 200 µM, but not 50 µM, transiently alleviated CCI-induced mechanical hypersensitivity assayed by von Frey test in male WT mice (**Fig. 7F-7H**), suggesting the potential of this clinically available VRAC inhibitor for neuropathic pain treatment. Treating *Swell1* cKO mice with dicumarol did not generate additional effects, indicating selectivity (**Fig. S7**). Furthermore, treating naïve mice with dicumarol did not change basal pain sensitivity and motor function (**Fig. S8**).

## DISCUSSION

Osmotic swelling of various cell types leads to a VRAC-dependent increase in extracellular ATP concentration (*41*). However, direct evidence establishing VRAC as an ATP-permeating channel is lacking. Moreover, the functional significance of VRAC-mediated ATP release in physiology and disease conditions remains unclear. In the present study, we provide electrophysiological evidence that Swell1-containing VRAC directly conducts and releases ATP. Functionally, the Swell1 channel in microglia contributes to nerve injury-induced increase in extracellular ATP content within the spinal cord, and plays an important role in spinal microglial activation, dorsal horn neuronal hyperactivity, and neuropathic pain-like behaviors. We further identify two FDA-approved drugs (dicumarol and zafirlukast) as novel and potent VRAC inhibitors and provide proof-of-concept for the potential translation of spinal Swell1-targeted therapies for the treatment of neuropathic pain by demonstrating the alleviation of mechanical allodynia after CCI with intrathecal administration of dicumarol in mice.

In addition to microglia in the spinal cord, the Cx3cr1-Cre line used in this work also drives Cre-mediated gene deletion in brain microglia and peripheral macrophages (*25*), both of which are also implicated in neuropathic pain (*42*). Although we cannot exclude the potential contribution of VRAC deletion in brain microglia and peripheral macrophages to *Swell1* cKO phenotype, our pharmacological data complement the genetic study and suggest that Swell1 channel in the spinal microglia is critical for the pathogenesis of neuropathic pain. Specifically, incubating spinal cord slices with dicumarol *ex vivo* dampens CCI-induced abnormal synaptic transmission of spinal dorsal horn neurons. Furthermore, intrathecal administration of dicumarol reversed CCI-induced mechanical hypersensitivity.

How does VRAC in the spinal microglia regulate neuropathic pain? Since microglia respond rapidly to peripheral nerve injury, we speculate that Swell1 channel in microglia is selectively activated by various inflammatory signals, including S1P. Swell1-mediated ATP release further activates microglial and neuronal P2X and P2Y receptors. This autocrine-paracrine purinergic signaling drives microglial activation, leading to the secretion of neuroactive factors (e.g. IL-1β) and aberrant activity of dorsal horn neurons. Future investigations using gene expression profiling and live-cell imaging will shed more light on the detailed mechanisms underlying Swell1 channel-mediated microglia activation. Notably, using a similar cKO mouse model, a very recent study reported that loss of Swell1 did not affect morphological changes of brain microglia upon ischemic stroke injury (*43*). In addition, Swell1 appears to be dispensable for microglia migration and chemotaxis following focal laser injury in the brain (*43*). Thus, the regulation of microglial activation by Swell1 may be dependent on the injury type, location, and specific pathophysiological context.

Given the importance of purinergic signaling in pain, several other ATP-releasing mechanisms have also been implicated in neuropathic pain conditions. Both genetic and pharmacological studies showed that connexin-43 and pannexin-1, two other large-pore channels (*44*), contribute to neuropathic pain (*16, 45, 46*), although it is yet to be determined if they regulate extracellular ATP concentration in the spinal cord. Moreover, exocytosis of ATP-containing vesicles from spinal dorsal horn neurons is critical for extracellular ATP enhancement and pain hypersensitivity following nerve injury (*24*). Interestingly, this mechanism does not appear to involve microglia activation as mice lacking vesicular nucleotide transporter (VNUT), a secretory vesicle protein responsible for the packaging and release of ATP, exhibited similar morphological changes and proliferation of spinal microglia as WT controls (*24*). There are also reports of potential off-target gene deletion in neurons for the Cx3cr1-Cre line (*47*), although this may be partially due to leaky reporter genes (*48*). Nevertheless, we cannot exclude the possibility that Swell1 channel in neurons is also involved in ATP release and pain hypersensitivity. Future work is necessary to reveal how the different ATP release mechanisms and cell types collectively regulate extracellular ATP levels in the spinal cord, thereby inducing neuropathic pain.

VRAC is a ubiquitously expressed ion channel, and its currents have also been observed in other glial cells, including astrocytes (*19*). However, deleting Swell1 and abolishing VRAC activity in microglia alone significantly reduced CCI-induced neuropathic pain-like behaviors in male mice. This highlights the importance of microglia and neuron-microglia interactions in neuropathic pain, an emerging theme from the recent literature in the field. Nevertheless, it is important to note that the Swell1 channel in brain astrocytes mediates glutamate release and promotes ischemic brain damage through excitotoxicity (*19, 49*). Thus, the alleviation of mechanical hypersensitivity by acute dicumarol treatment may not be mediated by inhibiting microglial VRAC and purinergic signaling alone. Future studies involving selective deletion of Swell1 in spinal astrocytes will reveal the role of neuron-astrocyte interaction and glutamate in the development and maintenance of neuropathic pain.

Most VRAC inhibitors currently available have low potency and poor selectivity (*34*). While DCPIB is a relatively potent and widely-used VRAC inhibitor, it is rather unspecific and toxic to cells, inhibiting or activating half a dozen other ion channels and transporters (*50–53*). Our identification of dicumarol and zafirlukast as potent and selective VRAC blockers provides much-needed pharmacological tools to not only probe the function of Swell1 channel, but also evaluate its therapeutic potential. Importantly, the achievable plasma level of dicumarol is significantly higher than its IC_50_ value for inhibition of VRAC activity, creating a wide therapeutic window for repurposing dicumarol as a novel treatment of neuropathic pain. Indeed, we show that intrathecal delivery of dicumarol acutely alleviates CCI-induced mechanical allodynia. Future work using different drug treatment protocols and additional outcome measures (e.g., spontaneous and ongoing pain) and pain models will further establish the analgesic profiles of Swell1-inhibiting drugs. Together with other recent screens (*37, 54*), our findings provide two novel drug scaffolds for further improving their potency and pharmacological properties (e.g. blood-brain barrier permeability). This will pave the way for therapeutic translation of better Swell1-targeting therapies for neuropathic pain and other diseases associated with abnormal VRAC activity, such as ischemic stroke.

## MATERIALS AND METHODS

### Mice

The protocol for all mice used in this study was approved by the Johns Hopkins University Animal Care and Use Committee. All experimental procedures involving vertebrate animals were conducted in accordance with the approved protocol MO22M17 and MO22M306. All mice were group-housed in a standard 12 hours light/ 12 hours dark cycle with ad libitum access to food and water. Both sexes were used in this study and specified in different assays. *Swell1^F/F^* mice were generated in the lab as previously described (*19*), and *Cx3cr1-Cre* (B6J.B6N(Cg)-Cx3cr1^tm1.1(cre)Jung/J^) were purchased from The Jackson Laboratory (Stock #025524). Adult C57BL6/J mice (2-4 months) mice were used for all animal related experiments. We used both male and female mice for our experiments and used age- and sex-matched littermates randomly assigned to different experimental groups.

### Chemicals

Chemicals are commercially available: DCPIB (Sigma-Aldrich, # SML2692-25MG), ARL 67156 trisodium salt (Tocris, #1283), sphingosine-1-phosphate (Cayman, # 62570), W146 (TOCRIS, # 36024), JTE013 (TOCRIS, # 2392), BAPTA AM (Cayman, # 15551), diphenyleneiodonium (DPI) (TOCRIS, # 0504), ATP (Sigma-Aldrich, # A2382-5G), LPS (Sigma-Aldrich, # L2880-100MG), nigericin (Cayman, 11437), dicumarol (Acros Organics, # 204120050), chenodiol (VWR 21947-5G), tioxolone (Sigma-Aldrich, # 217077), tolfenamic acid (Sigma-Aldrich, # T0535-5G), flufenamic acid (Acros Organics, # 165920100), zafirlukast (TCI America, #Z0029-100MG), 4-hydroxycoumarin (Acros Organics, # 121100050) and warfarin (Agilent, #PST-1000). All chemicals were dissolved with manufactures’ instructions. Dicumarol stock solution (10-20 mM) was made in dimethyl sulfoxide (DMSO) with brief sonication (2-5min) and heating (50 °C).

### Cell culture

HeLa/HEK293T/BV2 cell lines were maintained in Dulbecco’s modified Eagle’s medium (DMEM) supplemented with 10% fetal bovine serum (FBS) and 1% penicillin/streptomycin (P/S) at 37°C in humidified 95% CO_2_ incubator. For sniffer patch recordings, HeLa cells were digested in 0.25% trypsin and plated onto Poly-D-lysine coated 12-mm coverslips 1 d before the experiments. HEK293T cells were transfected with P2X2R-YFP (Addgene plasmid #22400) by Lipofectamine 2000 (Invitrogen) following the manufacturer’s instructions. On the day of sniffer patch, HEK293T cells expressing P2X2R-YFP were dissociated, triturated, and added onto the coverslips with HeLa cells. *SWELL1* knockout (KO) HeLa cells were reported previously(*55*).

*SWELL1* knockout (KO) BV2 microglia were generated using CRISPR/Cas9 technology. Guide RNA (GCTGTGTGTCCGCAAAGTAG) targeting murine LRRC8A (Swell1) was cloned into LentiCRISPR-v2-PuroR, which was a gift from Feng Zhang (Addgene plasmid #98290). The primers used to design the sgRNA targets were (5’ to 3’) Swell1 forward caccGCTGTGTGTCCGCAAAGTAG; Swell1 reverse aaacCTACTTTGCGGACACACAGC. Lentiviral particles containing Swell1 sgRNA were packaged using the 3rd generation lentiviral system and used to infect BV2 microglia. One day post-infection, the medium was changed to fresh DMEM containing 10% FBS and 1% PS. Cells were then treated with 5 mg/ml puromycin for 5 days to select for successfully transduced cells. Single clones were obtained using limiting dilution and were characterized using genotyping analysis of frameshift mutations by target-site-specific PCR and TA cloning followed by sanger sequencing. Swell1 protein expression was validated in single clones using western blotting.

Bone marrow-derived macrophages (BMDMs) were freshly isolated and cultured from adult C57BL6/J mice (2-4 months). Bone marrow was flushed from both femurs using a syringe and 26-gauge needle, and briefly centrifuged to get rid of tissue chunks. BMDMs were cultured and differentiated in 70% Roswell Park Memorial Institute (RPMI) 1640 Medium (20% FBS, 1% penicillin/streptomycin (P/S)) supplemented with 30% L929 mouse fibroblast-conditioned media for 6–7 days. For electrophysiology recording and IL-1β detection, BMDMs were scraped and reseeded 1 day before the experiments.

### Immunostaining

Following procedures previously described in (*19*), we perfused anesthetized mice transcardially with phosphate•buffered saline (PBS), followed by 4% paraformaldehyde (PFA) in PBS. Spinal cords were removed and post-fixed in 4% PFA at 4 °C overnight. After dehydration by 30% sucrose, spinal cords were embedded in OCT (Tissue-Tek) and cut into 35-μm-thick sections on a cryostat microtome (Leica). Sections were permeabilized with 0.2% Triton X-100 and 1% BSA in PBS for 45 min at room temperature (RT), washed with PBS three times, blocked in 10% BSA, and incubated with primary antibodies at 4 °C overnight. Primary antibody concentrations: anti-Iba1 (1:500; Wako), anti-cFos (1:500; Millipore), anti-NeuN (1:500; Millipore). After washing with PBS 3 times, samples were incubated with Alexa Fluor-conjugated secondary antibodies (1:500; Invitrogen) for 1 h at RT. Z-stack images were taken under Zeiss LSM800 or 900 confocal microscopes.

Z-stack images were orthogonal projected before the analysis. To determine the intensity of Iba1, similar areas of spinal cord dorsal horn was selected from each section and the average intensities were measured by ImageJ. The cell number was quantified manually with ImageJ plugins Cell Counter (NIH) as previously described (*56*). To examine the c-Fos positive neurons, positive cell numbers in the spinal cord dorsal horn were counted using ImageJ built-in ‘Analyze Particles’ function with manual inspection. All analyses were performed by investigators blinded to mouse genotype.

### RNAscope *in situ* hybridization

Fixed brains were embedded in OCT (Tissue-Tek) and sectioned at a thickness of 14 µm. RNAscope Multiplex Fluorescent Reagent Kit v2 (ACD, Advanced Cell Diagnostics) was used following the manufacturer’s manual for the fixed frozen tissues (*19*). Probe targeting Swell1 (#458371) was purchased from ACD. TSA Plus fluorescein (#NEL741E001KT) was used for developing the fluorescence signal. After *in situ* hybridization, sections were subjected to Iba1 immunostaining as described above. Images were collected under a Zeiss LSM 900 confocal microscope.

### Quantitative real-time PCR

For the analysis of the S1P receptor expression profile in BV2 cells, total RNA was isolated using TRIzol reagent (Life Technologies). 2 µg of total RNA was used to generate the cDNA using the High-Capacity cDNA Reverse Transcription Kit (ThermoFisher Scientific). The reaction was run in QuantStudio 6 Real-Time PCR Systems using 0.2 μl of cDNA in a 15 μl reaction according to the manufacturer’s instruction in triplicate that measures real-time SYBR green fluorescence. Calibrations and normalizations were performed using the 2^-ΔΔCT^ method. *Gapdh* was used as the reference gene.

Designed mouse *Gapdh* assay (forward primer: AACTTTGGCATTGTGGAAGG; reverse primer: GGATGCAGGGATGATGTTC). *S1pr1*: (forward primer: ACTACACAACGGGAGCAACAG, reverse primer: GATGGAAAGCAGGAGCAGAG). *S1pr2*: (forward primer: CTCACTGCTCAATCCTGTCATC, reverse primer: TTCACATTTTCCCTTCAGACC). *S1pr3*: (forward primer: TTCCCGACTGCTCTACCATC, reverse primer: CCAACAGGCAATGAACACAC). *S1pr4*: (forward primer: TGCGGGTGGCTGAGAGTG, reverse primer: TAGGATGAGGGCGAAGACC). *S1pr5*: (forward primer: CTTAGGACGCCTGGAAACC, reverse primer: CCCGCACCTGACAGTAAATC).

### Western blot

Homogenates of cultured BV2 microglia were prepared in lysis buffer containing (in mM): 20 Tris-HCl, pH 7.5, 150 NaCl, 1% triton and 1% protease inhibitors cocktails. Samples were resolved on SDS/PAGE and transferred to nitrocellulose membranes, which were incubated in the Tris-buffered saline (TBS) buffer containing 0.1% Tween-20 and 5% milk for 1 h at RT before the addition of primary antibody (anti-Swell1; 1:1000; from a rabbit immunized with Swell1 C-terminal peptide antigen) for incubation overnight at 4 °C (*19*). After wash, the membranes were incubated with HRP-conjugated secondary antibody (Thermo Scientific) in the same TBS buffer for 1 h at RT. Immunoreactive bands were visualized using enhanced chemiluminescence. Films were scanned with a scanner and analyzed with Image J.

### IL-1β ELISA

For IL-1β release from BMDMs, cells were primed with LPS (1 µg/mL, 4 h) in DMEM (10% FBS, 1% PS). After priming, the media was replaced with pure serum-free DMEM or serum-free DMEM with 5 mM ATP or 10 µM nigericin, or when specified an isotonic buffer (in mM): (140 NaCl, 5 KCl, 1 MgCl_2_, 1 CaCl_2_, 10 HEPES, 5 glucose, pH 7.3, 310 mOsm/kg) or hypotonic buffer (40 NaCl, 5 KCl, 1 MgCl_2_, 1 CaCl_2_, 10 HEPES, 5 glucose, pH 7.3, 110 mOsm/kg). After 1-4 h, supernatants were collected and the released IL-1β were measured by mouse IL-1β/IL-1F2 DuoSet ELISA (Cat. number: DY401) from R&D Systems.

For spinal cord IL-1β detection, the L3–L5 lumbar spinal cord segment was removed 7 days after CCI surgery and rapidly immersed in RIPA lysis buffer with 1% protease inhibitors cocktails. Total IL-1β protein level was measured by mouse IL-1β/IL-1F2 DuoSet ELISA kit and first normalized to total protein level quantified by Pierce™ BCA Protein Assay Kit (Thermo Scientific), and then normalized to the sham group.

### ATP measurement

HeLa and BV2 cells cultured in 24-well plates were grown to 70–80% confluence. Before measurement, the cell culture medium was replaced by isotonic buffer (in mM): (140 NaCl, 5 KCl, 1 MgCl_2_, 1 CaCl_2_, 10 HEPES, 5 Glucose, pH 7.3, 310 mOsm/kg) to reduce the ATP background. Cells were treated with hypotonic buffer (70 NaCl, 2.5 KCl, 0.5 MgCl_2_, 0.5 CaCl_2_, 5 HEPES, 2.5 Glucose, pH 7.3, 155 mOsm/kg) or 10 nM S1P in isotonic buffer to induce VRAC activation. Different concentrations of DCPIB/dicumarol were added without pretreatment in specified conditions. The supernatant was collected after 30 min of treatment for the luciferase assay. For BV2 cells, brief centrifugation was used to get rid of floating cells. ATP concentration was measured using a luciferin–luciferase ENLITEN® ATP Assay System (Promega). Luminescence was measured by Infinite M Plex microplate reader (Tecan).

For extracellular ATP measurement in the spinal cord (*24*), the L3–5 spinal cord was removed immediately following spinal laminectomy. Acute spinal cord slices were incubated in ice-cold artificial cerebrospinal fluid (ACSF) containing (in mM): 125 NaCl, 2.5 KCl, 2.5 CaCl_2_, 1.3 MgCl_2_, 1.3 NaH_2_PO_4_, 26 NaHCO_3_, 10 glucose, saturated with 95% O_2_ and 5% CO_2_ with the ectonuclease inhibitor ARL67156 (100 μM) (Tocris) for 20 min. Extracellular ATP levels were measured from the ACSF with the same ATP assay system and plate reader mentioned above and normalized to the total protein level measured by Pierce™ BCA Protein Assay Kit.

### Electrophysiology

For hypotonicity-activated VRAC current recordings (*19*), whole-cell patch-clamp configuration was established in an isotonic bath solution containing (in mM): 90 NaCl, 2 KCl, 1 MgCl_2_, 2 CaCl_2_, 10 HEPES, 10 glucose, 100 mannitol (pH adjusted to pH 7.3 with NaOH and osmolality adjusted to 310 mOsm/kg), then hypotonic solution which has the same ionic composition but without mannitol were applied. Recording electrodes (2-4 MΩ) were filled with a standard internal solution containing (mM):133 CsCl, 10 HEPES, 4 Mg-ATP, 0.5 Na_3_-GTP, 2 CaCl_2_, 5 EGTA (pH adjusted to 7.2 with CsOH and osmolality was 290-300 mOsm/kg). For S1P activated VRAC current recordings, the bath solution contained (in mM): 145 NaCl, 2 KCl, 1 MgCl_2_, 2 CaCl_2_, 10 HEPES, 10 glucose (pH adjusted to pH 7.3 with NaOH and osmolality adjusted to 300-310 mOsm/kg). For the VRAC time course, constant voltage ramps (5 s interval, 500 ms duration) were applied from a holding potential of 0 to ± 100 mV. For the step protocol, cells were held at −60 mV and voltage step pulses (3 s interval, 500 ms duration) were applied from −100 to +100 mV in 20 mV increments. To determine the ATP permeability of VRAC, recording electrodes (2-4 MΩ) were filled with the Cl^−^ based intracellular solution containing (in mM): 50 NaCl, 10 HEPES, 200 mannitol (pH adjusted to pH 7.3 with NaOH and osmolality adjusted to 310 mOsm/kg). ATP^4−^ based intracellular solution was made by replacing 50 NaCl and 200 mannitol with 50 Na_2_ATP^2−^ and 150 mannitol to maintain the same osmolality. The reversal potentials were determined using the ramp protocol. Relative permeability of ATP^4−^ to Cl^−^ was estimated based on the Goldman– Hodgkin–Katz flux equation and solved by the Newton Raphson technique(*57*).

For sniffer patch recordings (*19*), HeLa cells were used as the source cells, and HEK293T cells transfected with P2X2R-YFP were the sensor cells. 12-18 h after transfection, HEK293T cells were reseeded onto source cells for recordings. To activate VRAC in HeLa cells, recording electrodes (2-4 MΩ) were filled with a hypertonic internal solution containing (mM): 133 CsCl, 10 HEPES, 10 Mg-ATP, 0.5 Na_3_-GTP, 2 CaCl_2_, 5 EGTA, 100 mannitol (pH adjusted to 7.2 with CsOH and osmolality was 400-410 mOsm/kg). For HEK293T cells recording, the same internal solution was used except there was no mannitol. External solution contained (in mM): 145 NaCl, 10 HEPES, 2 KCl, 2 CaCl_2_, 1 MgCl_2_, and 10 glucose (pH adjusted to pH 7.3 with NaOH and osmolality adjusted to 300-310 mOsm/kg). HeLa cells were held at −100 mV, and HEK293T cells were held at −60 mV. Recordings were made with MultiClamp 700B amplifier and 1550B digitizer (Molecular Device). Data acquisitions were performed with pClamp 10.7 software (Molecular Device), filtered at 1 kHz and digitized at 10 kHz.

### Acute spinal cord slice electrophysiology

To prepare and record acute spinal cord slices (*19*), mice (2-3 months old) were anesthetized with the inhalation of anesthetic isoflurane, and then perfused with the ice-cold oxygenated cutting solution. The L3–L5 lumbar spinal cord segment was removed rapidly and immersed in ice-cold choline-based cutting solution containing (in mM): 110 choline chloride, 7 MgCl_2_, 2.5 KCl, 0.5 CaCl_2_, 1.3 NaH_2_PO_4_, 25 NaHCO_3_, 20 glucose, saturated with 95% O_2_ and 5% CO_2_. Coronal spinal cord slices (400 µm) were cut in the cutting solution using a vibratome (VT-1200S, Leica) and transferred to artificial cerebrospinal fluid (aCSF) containing (in mM): 125 NaCl, 2.5 KCl, 2.5 CaCl_2_, 1.3 MgCl_2_, 1.3 NaH_2_PO_4_, 26 NaHCO_3_, 10 glucose, saturated with 95% O_2_ and 5% CO_2_. The slices were allowed to recover for 1 h at 32 °C and then at RT for at least 1 h before recording. All recordings were made at RT in a submerged recording chamber with constant ACSF perfusion. Whole-cell recordings from lamina II neurons were visualized under an upright microscope (BX51WI, Olympus) with infrared optics. Recording pipettes (3-5 MΩ) were filled with an internal solution containing (in mM): 125 K-gluconate, 15 KCl, 10 HEPES, 1 MgCl_2_, 4 Mg-ATP, 0.3 Na_3_-GTP, 10 phosphocreatine, and 0.2 EGTA (pH 7.2, osmolality 290-300 mOsm/kg). To block inhibitory synaptic transmission, picrotoxin (100 µM) was added to the bath in all recordings. For sEPSC recordings, cells were held at −70 mV. In pharmacological experiments, 10 min before recording, dicumarol (20 µM) was bath applied. Recordings were made with MultiClamp 700B amplifier and 1550B digitizer (Molecular Device). Data acquisitions were performed with pClamp 10.7 software (Molecular Device), filtered at 1 kHz and digitized at 10 kHz. In all experiments, the series resistance (Rs) was monitored throughout the recording and controlled below 20 MΩ with no compensation. Data were discarded when the series resistance varied by ≥ 20%.

### Behavioral study

All behavioral assays were conducted by an observer who was blinded to genotype or drug condition. Mature mice were used for all behavior assays (2–4 month). Mice were habituated before any behavioral test.

#### Open field test

Open field test was used to assess the locomotor activity of the mice. Mice were placed in an open field chamber (73 × 45 cm rectangular plastic box with a wall height of 33 cm) for 10 min. We analyzed parameters such as total distance traveled in video recordings using SMART 3 software (Panlab Harvard Apparatus, Barcelona, Spain).

#### Mechanical paw withdrawal frequency test

To measure paw withdrawal responses to repeated mechanical stimuli, mice were placed in individual Plexiglas cages with a wire mesh floor and allowed to acclimate for 30min. Two calibrated von Frey monofilaments (0.07 and 0.4 g) were used. Each von Frey filament was applied perpendicularly to the plantar side of the hind paw for approximately 1 s, and each stimulation was repeated 10 times to both hind paws. The occurrence of paw withdrawal in each of these 10 trials was expressed as a percent response frequency [(number of paw withdrawals/10 trials) × 100 = % response frequency], and this percentage was used as an indication of the amount of paw withdrawal and mechanical sensitivity.

#### Hargreaves test

To test for signs of heat hypersensitivity, we used the Hargreaves test(*58*) which measures paw-withdrawal latency (PWL) to radiant heat stimuli. Animals were placed under individual transparent plastic boxes on a heated glass floor (30 °C) and allowed to habituate for at least 30 min before testing. Radiant heat was applied to the plantar surface of each hind paw three times at 3–5-min intervals with a plantar stimulator analgesia meter (IITC model 390, Woodland Hills, CA). PWLs were measured three times by an electronic timer, with at least 2 min between trials. A cut-off time of 30 s was used to avoid sensitization and damage to the skin. The average PWL of three trials was used for data analysis.

#### Catwalk gait analysis

Animals were subjected to gait analysis using the CatWalk Automated Gait Analysis System (Noldus Information Technology, Leesburg, VA) and software version XT 10.6. Gait analysis was performed in mice basally as well as 7-11 days after CCI surgery. Each animal completed three runs (n = 3). Subjects traverse a walkway (31.7 cm long; 6.36 cm wide) atop a glass floor in a darkened room. Light enters the distal long edge of the glass floor from a fluorescent bulb located at the side, and is internally reflected, scattering only at points where a paw touches the glass, producing bright illumination of the contact area. Internal reflection is incomplete, allowing a faint superimposed image of the animal to be seen as well. A video camera monitors the corridor, and the digitized signal is stored for later analysis of animals crossing the walkway. Paws were first auto classified followed by manual inspection. Parameters related to pain behaviors and motor functions were analyzed. Average speed is the speed of the animal’s body in the recorded run. Max Contact Area is the maximum area of a paw that comes into contact with the glass plate. Max Intensity is the maximum intensity of the complete paw. Stand is the duration of contact of a paw with the glass plate. Swing is the duration of no contact of a paw with the glass plate. Duty Cycle (%) expresses Stand as a percentage of Step Cycle: Stand/ (Stand + Swing) × 100%.

### Chronic constriction injury model of neuropathic pain

The chronic constriction injury was performed as previously described (*26*). Three loose nylon monofilament (Covidien Monosof, 9-0, #N2533) ligatures were put around the unilateral sciatic nerve. The incision was closed with sutures or tissue glue. Sham-operated mice received only an identical dissection of the unilateral nerve exposure and nerve isolation without the ligature. Behavioral tests were performed 1, 3, 5, 7, and 11 days after peripheral nerve injury.

### Intrathecal drug administration

#### Hypotonicity-induced YFP quenching assay

HEK293T cells stably express halide-sensitive YFP was generated as previously described (*17*). Cells were seeded in 384 well plate with 90% confluence one night before the assay. Cells were washed three times with isotonic buffer (in mM): (140 NaCl, 2 KCl, 1 MgCl_2_, 1 CaCl_2_ 10 HEPES, pH adjusted with NaOH to 7.25-7.3, osmolality 290-300 mOsm/kg.) to reduce basal fluorescence from culture media. We stimulated cells with 15 μL/well hypotonic solutions (in mM): (5 KCl, 20 HEPES, 110 mM Mannitol) for 5 min, followed by addition of 10 μL/well 75 mM NaI. For drug screening, 5 ul of stock compound solution (final concentration 20 µM) was added to each well after cell wash and incubated for 10 min at room temperature before moving to FDSS6000. FDSS6000 (Hamamatsu Photonics) recorded fluorescence in each well simultaneously during the assay. The fluorescence reading after NaI addition was normalized to that at 1 s before the addition (set as 100%). Z factor is estimated from the sample means and sample standard deviations of positive (WT cells) and negative (*SWELL1* KO cells) controls. Z score, also known as the standard score, is determined by the number of standard deviations by which the value of raw data is above or below the mean value.

#### Statistical analysis

Following procedures previously described in (*19*), we used GraphPad Prism 9.1 software for all statistical analyses. Images were analyzed with ImageJ (US National Institutes of Health) and ZEN software (ZEISS). All the behavior assays and image quantifications were performed by investigators blinded to mouse genotype. Before each test, the Gaussian distribution of the data was assessed using the Shapiro-Wilk normality test to determine whether the data were normally distributed. Parametric tests (paired or unpaired Student’s t-test for two groups) were used for normally distributed data. ANOVA followed by Bonferroni’s post hoc test was used to analyze results with more than two groups. Data are reported as mean ± SEM. The significance level was set at p < 0.05. The number (n) of cells and animals for each experiment are shown in each figure and described in each figure legend.

## Acknowledgements

We thank H. Xu and A. Long at Johns Hopkins University (JHU) High Throughput Screening ChemCORE for help with drug screening; Q. Zheng, and Q. Huang for help with mouse behavior; A. Kamiya for BV2 microglia cells; T. Connor for L929 cells; and X. Dong and M. Caterina for helpful discussion. This work was facilitated by the Pain Research Core, which is funded by the Blaustein Fund and the Neurosurgery Pain Research Institute at JHU. The Johns Hopkins Drug Library was supported by FAMRI, IBBS and the Kimmel Comprehensive Cancer Center of JHU (J.O.L.).

## Author Contributions

J.Chu designed and performed most of the experiments, and analyzed the data. J.Y. performed spinal cord slice electrophysiological recordings. Y.Z. performed the initial behavioral assays. K.H.C., J.Chen, H.Y.C., and N.K helped with the experiments. C.Z. assisted with the CatWalk assay. J.O.L. provided the Johns Hopkins Drug Library. Y.G. discussed the data, provided critical advice, and revised the manuscript. Z.Q. supervised the project. Z.Q., Y.G., and J.Chu wrote the manuscript with input from all authors.

## Data and materials availability

All data are available in the main text or the supplementary material.

---

**Fig. S1.**
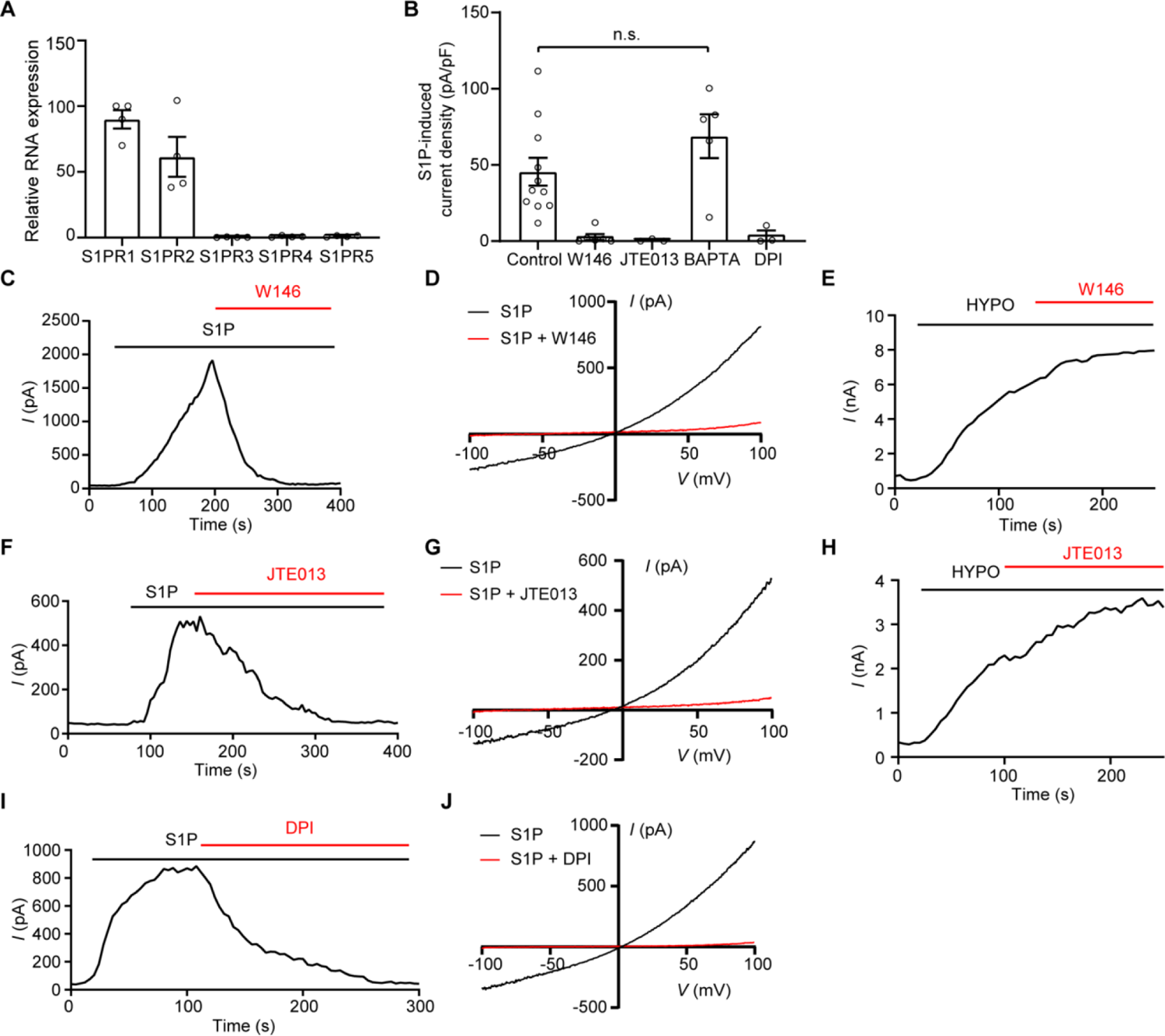
Characterization of the mechanisms underlying S1P-induced VRAC activation in BV2 microglia. (A) S1PRs expression in BV2 microglia assayed by qPCR. Gapdh was used as the reference gene. n = 4 biological replicates. (B) Quantification of 10 nM S1P-induced VRAC current densities at +100 mV in BV2 microglia with no treatment (control; n = 11 cells) or treated with 10 μM W146 (selective S1PR1 antagonist; n = 6 cells), 100 nM JTE013 (selective S1PR2 antagonist; n = 3 cells), 5 mM BAPTA (Ca^2+^ chelator; n = 5 cells), 20 μM DPI (NAD(P)H oxidase inhibitor; n = 3 cells). S1P-induced VRAC currents in microglia are sensitive to inhibitors of S1PR1, S1PR2 and ROS production, but not Ca^2+^ chelation. One-way ANOVA, Bonferroni post hoc test, n.s.: not significant. (**C-D**) Representative time course (C) at +100 mV and I-V curve (D) of S1P-induced VRAC currents treated with 10 µM W146. (**E**) Representative time course at +100 mV of hypotonicity (HYPO; 200 mOsm/kg)-induced VRAC currents in microglia treated with 10 µM W146. W146 specifically inhibits S1P-, but not HYPO-induced VRAC activity. (**F-G**) Representative time course (F) at +100 mV and I-V curve (G) of S1P-induced VRAC currents treated with 100 nM JTE. (**H**) Representative time course at +100 mV of HYPO-induced VRAC currents in microglia treated with 100 nM JTE. JTE specifically inhibits S1P-, but not HYPO-induced VRAC activity. (**I-J**) Representative time course (I) at +100 mV and I-V curve (J) of S1P-induced VRAC currents treated with 20 µM DPI. Bar graphs are reported as mean ± SEM.

**Fig. S2.**
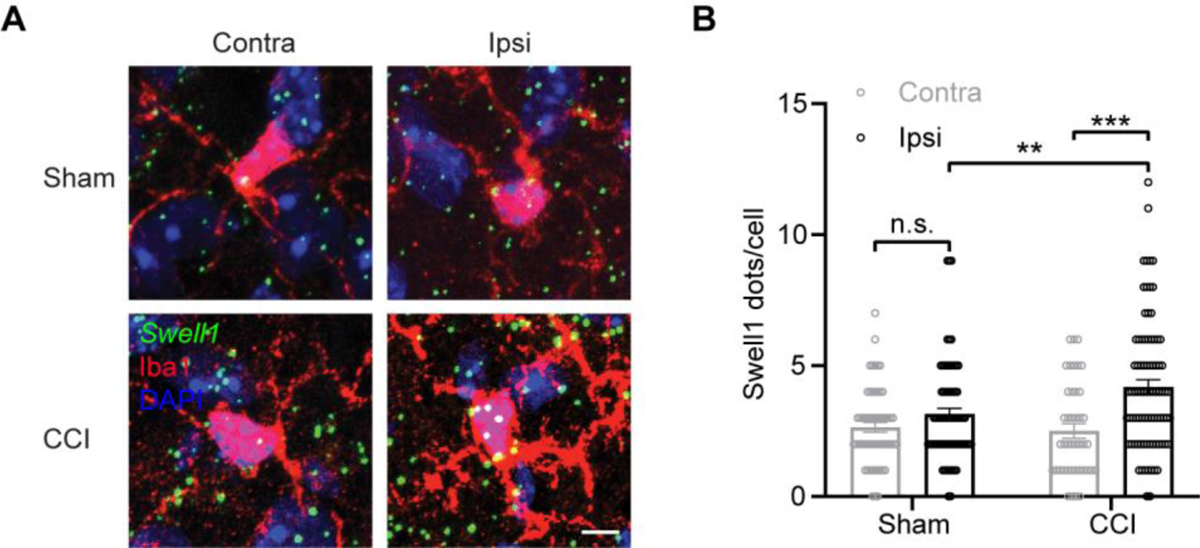
*Swell1* mRNA in microglia is upregulated after peripheral nerve injury. (**A**) Representative images and (**B**) quantifications (mean ± SEM) of *Swell1* RNAscope *in situ* hybridization and Iba1 co-staining in the spinal cord dorsal horn from ipsilateral and contralateral side of CCI or sham mice. n = 59 microglia cells for contralateral side, n = 75 microglia cells for ipsilateral side from 4 mice for sham group; n = 42 microglia cells for contralateral side, n = 89 cells for ipsilateral side from 4 mice for CCI group. Two-way ANOVA, Bonferroni post hoc test, n.s.: not significant, **p < 0.01, ***p < 0.001. Scale bar, 5 μm.

**Fig. S3.**
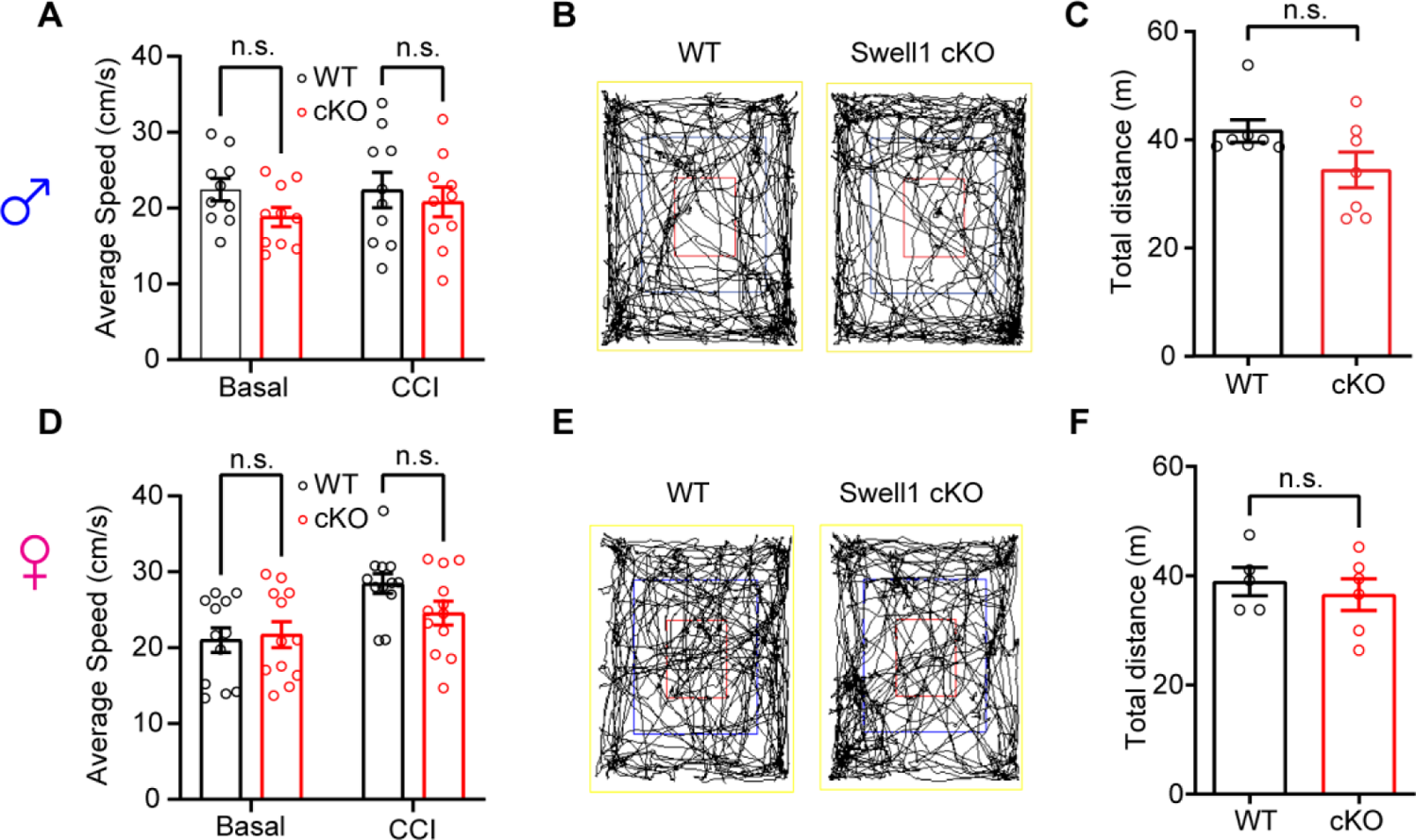
Microglia-specific *Swell1* cKO mice exhibit normal locomotor function in both sexes. (A) Quantification of average speed in CatWalk assay before and 7 days after CCI in male WT mice (n = 10) and *Swell1* cKO mice (n = 10). Two-way ANOVA, Bonferroni post hoc test, n.s.: not significant. (B) Representative traces of male mice during the open field test. (C) Quantification of total distance traveled in the open field test for male WT (n = 7) and *Swell1* cKO mice (n = 7). Student’s *t*-test, n.s.: not significant. (D) Quantification of average speed in CatWalk assay before and 7-11 days after CCI in female WT mice (n = 12) and *Swell1* cKO mice (n = 12). Two-way ANOVA, Bonferroni post hoc test, n.s.: not significant. (E) Representative traces of female mice during the open field test. (F) Quantification of total distance traveled in the open field test for male WT (n = 5) and *Swell1* cKO mice (n = 6). Student’s *t*-test, n.s.: not significant. Bar graphs are reported as mean ± SEM.

**Fig. S4.**
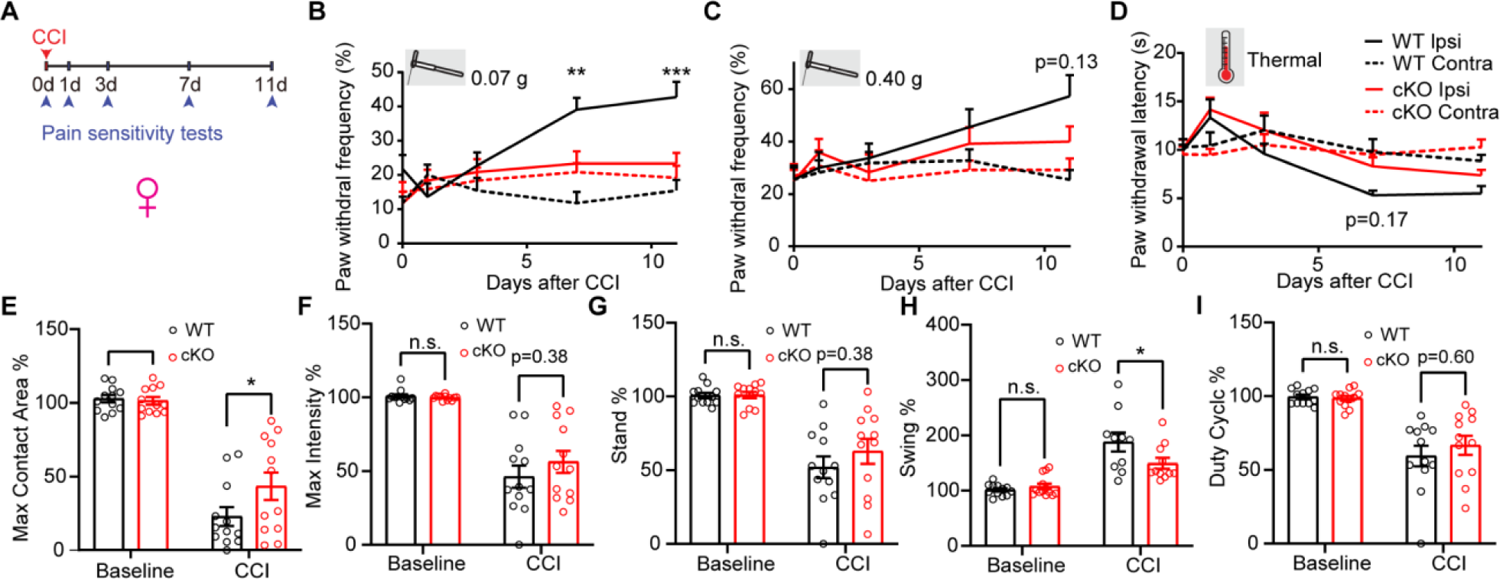
Characterization of CCI-induced neuropathic pain-like behaviors in female *Swell1* cKO mice. (**A**) Experimental design of measuring CCI-induced neuropathic pain-like behaviors. Baseline sensitivity was measured at day 0 before CCI. (**B-D**) Baseline and CCI-induced paw withdrawal frequency detected by the von Frey filaments (B: 0.07 g; C: 0.40 g) and paw withdrawal latency detected by Hargreaves test (D) in female WT (n = 11) and *Swell1* cKO (n = 11) mice from ipsilateral and contralateral sides. Statistics between WT ipsilateral side and *Swell1* cKO ipsilateral side were labeled. Two-way ANOVA, Bonferroni post hoc test, **p < 0.01, ***p < 0.001. **(E-I)** Quantification of the right hind paw (ipsilateral to the side of nerve injury) maximum contact area (E), intensity (F), stand (G), swing (H), duty cycle (I) normalized to left hind paw (contralateral to the side of nerve injury) at baseline and 7-11 days after CCI in female WT mice (n = 12) and *Swell1* cKO mice (n = 12). Two-way ANOVA, Bonferroni post hoc test, n.s.: not significant, *p < 0.05, **p < 0.01. Bar graphs are reported as mean ± SEM.

**Fig. S5.**
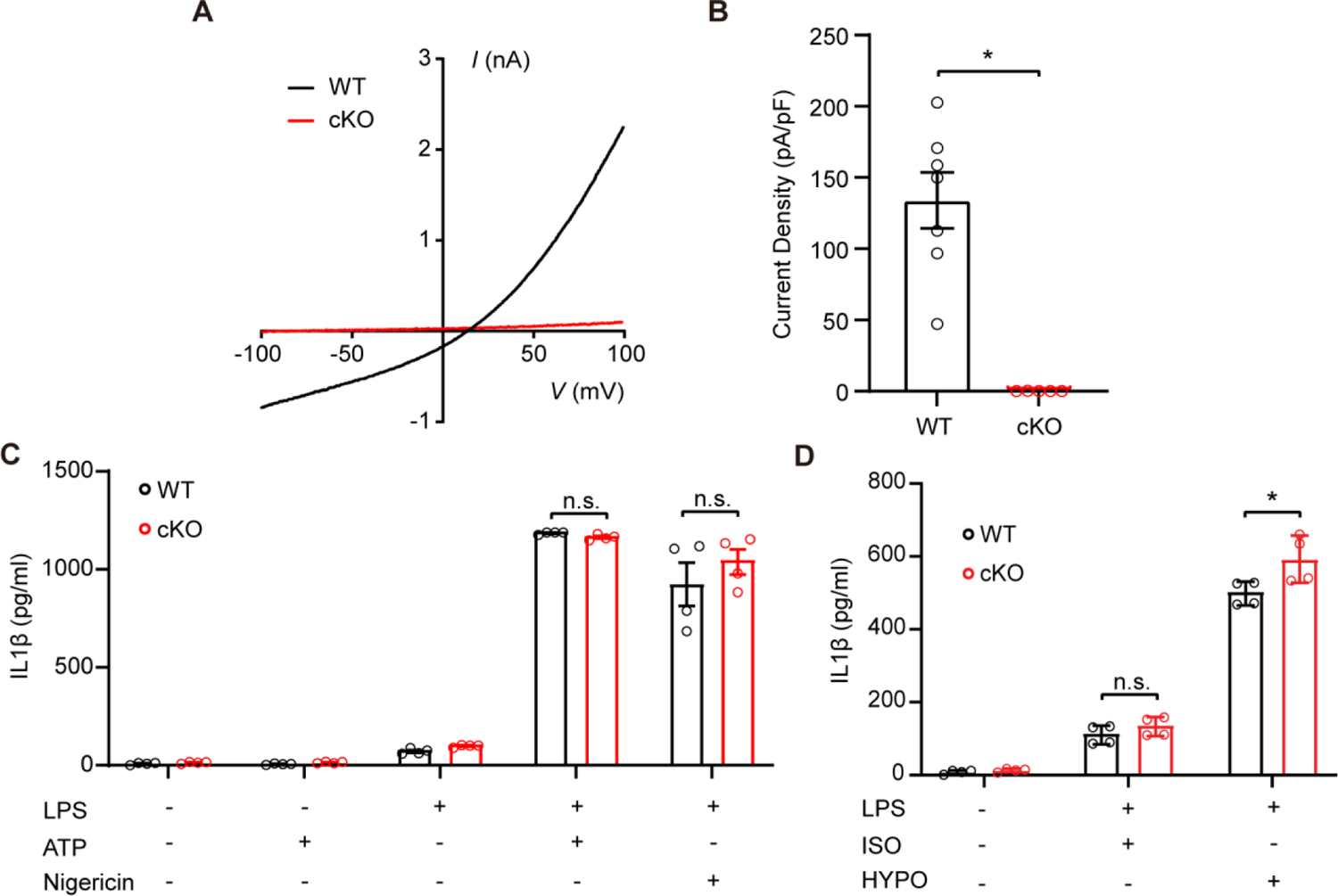
Swell1 is not required for DAMP- or hypotonicity-induced NLRP3 inflammasome activation. (A) Representative hypotonicity-induced VRAC currents recorded by voltage ramp protocols in BMDMs isolated from WT and *Swell1* cKO mice. (B) Quantification of hypotonicity-induced VRAC current densities at +100 mV in BMDMs isolated from WT and *Swell1* cKO mice. Student’s *t*-test, *p < 0.05. (C) IL-1β releases were determined by ELISA on supernatants in BMDMs isolated from WT and *Swell1* cKO mice. Naïve or LPS-primed (1 µg/mL for 4 h) BMDMs were treated with 5 mM ATP or 10 µM nigericin. Two-way ANOVA, Bonferroni post hoc test, n.s.: not significant. (D) IL-1β releases were determined by ELISA on supernatants in BMDMs isolated from WT and *Swell1* cKO mice. Naïve or LPS-primed (1 µg/mL for 4 h) BMDMs were stimulated with isotonic (300 mOsm/kg) or hypotonic solution (100 mOsm/kg) for 4 h. Two-way ANOVA, Bonferroni post hoc test, n.s.: not significant, *p < 0.05. Note that there is even a slight increase in IL-1β release from cKO BMDMs treated with LPS and HYPO compared WT cells. Bar graphs are reported as mean ± SEM.

**Fig. S6.**
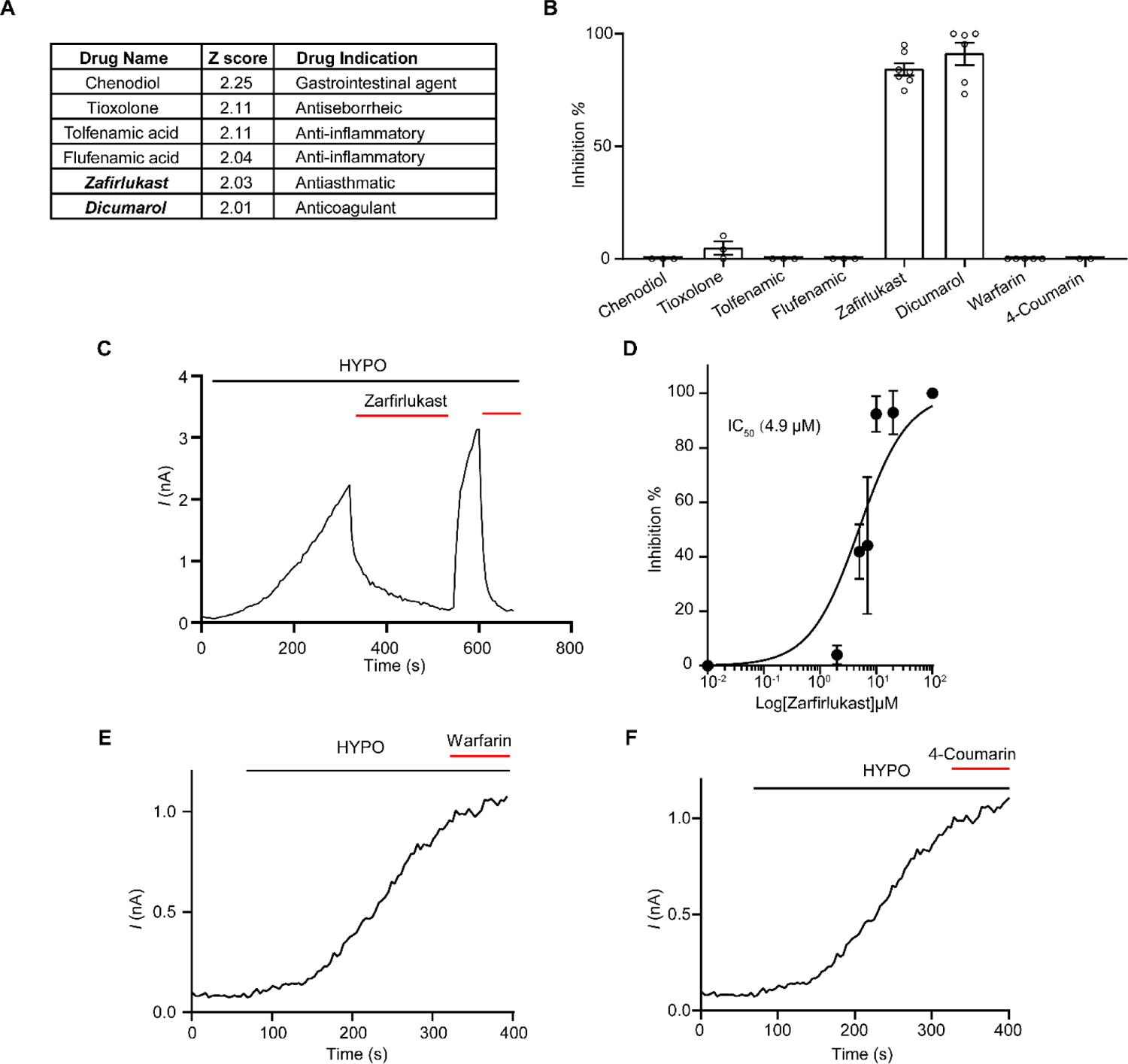
Characterization of VRAC inhibitor candidates from the high-throughput screening. (A) A list of VRAC inhibitor candidates from FDA-approved drug library. (B) Quantification of drug (20 µM)-mediated inhibition on hypotonicity-induced VRAC currents at +100 mV in HEK293T cells. (C) Hypotonicity-induced VRAC currents at +100 mV were inhibited by 20 µM Zafirlukast in HEK293T cells. HYPO: 200 mOsm/kg. (D) Dose-inhibition curve of zafirlukast on hypotonicity-induced VRAC currents in HEK293T cells (n = 3-9 cells). (**E-F**) HYPO-induced VRAC currents at +100 mV were not inhibited by 20 µM Warfarin (E) and 4-hydroxycoumarin (F). Bar graphs are reported as mean ± SEM.

**Fig. S7.**
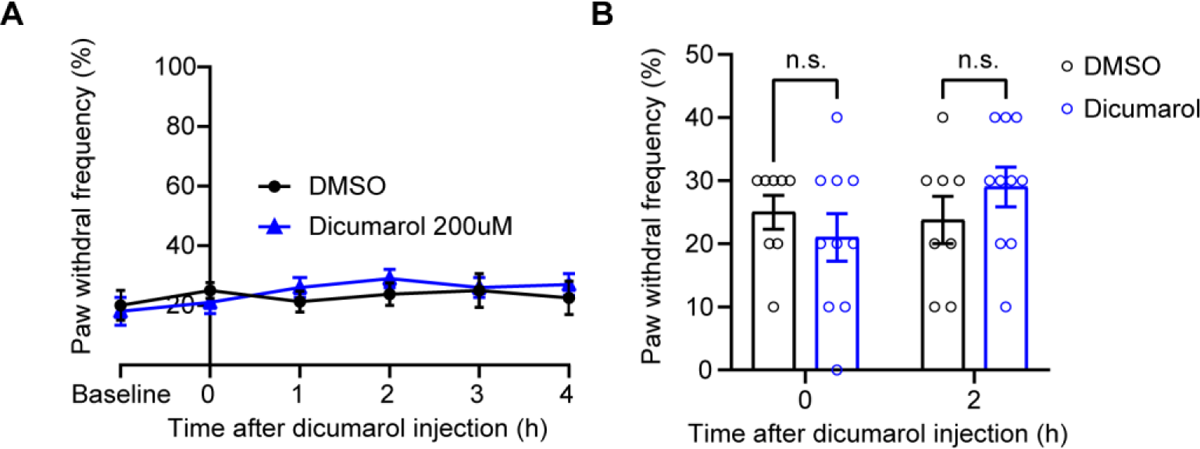
Dicumarol does not have any additional effects in microglia-specific *Swell1* cKO mice. (**A-B**) Paw withdrawal frequency was detected by the von Frey Filament (0.40 g) in CCI-treated *Swell1* cKO mice before and after intrathecal injection of DMSO or 200 µM dicumarol. Time course of the paw withdrawal frequency was plotted in (A), detailed bar graph before and 2 hours after the dicumarol injection was plotted in (B). n = 8 mice for DMSO, n = 9 mice for dicumarol. Two-way ANOVA, Bonferroni post hoc test, n.s.: not significant.

**Fig. S8.**
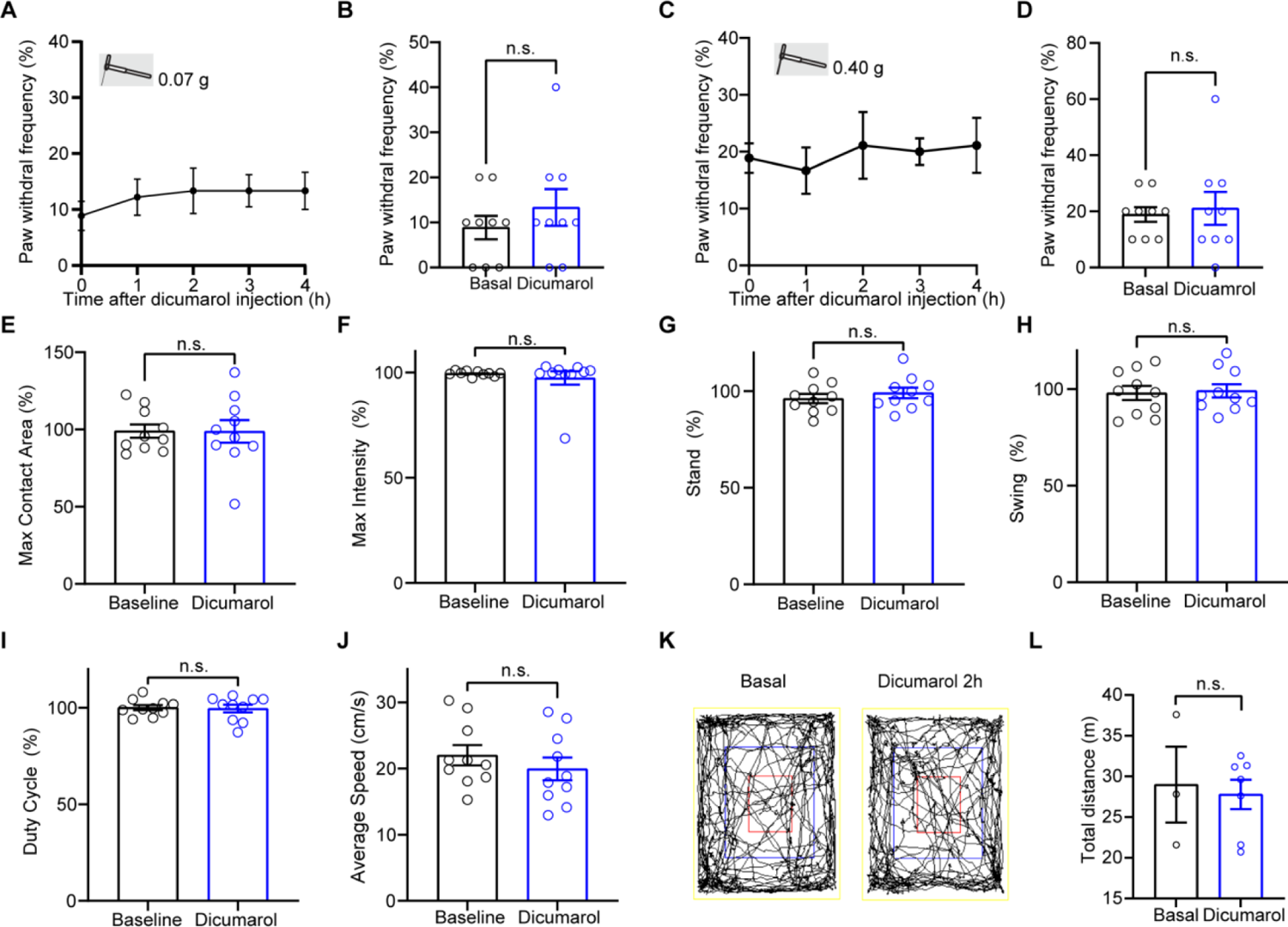
Dicumarol does not affect basal pain sensitivity and motor function in naïve mice. (**A**-**B**) Paw withdrawal frequency was detected by the von Frey Filament (0.07 g) WT mice before and after intrathecal injection of 200 µM dicumarol. Paw withdrawal frequency before injection (baseline) and 2 hours after injection(dicumarol) are plotted in (B). n = 9 mice. Student’s t-test, n.s.: not significant. (**C**-**D**) Paw withdrawal frequency was detected by the von Frey Filament (0.40 g) WT mice before and after intrathecal injection of 200 µM dicumarol. Paw withdrawal frequency before injection (baseline) and 2 hours after injection (dicumarol) are plotted in (D). n = 9 mice. Student’s t-test, n.s.: not significant. **(E-J)** Quantification of the maximum contact area (E), intensity (F), stand (G), swing (H), duty cycle (I), average speed (J), injection (baseline) and 2 hours after 200 µM dicumarol injection (dicumarol) in CatWalk assay. Right hind paw was normalized to left hind paw for paw specific statistics. n = 10 mice. Student’s t-test, n.s.: not significant. (**K**-**L**) Representative traces (K) and quantifications (L) of total distance traveled in the open field test for mice 2 hours after no injection (baseline) and 2 hours after 200 µM dicumarol injection (dicumarol). n = 3 mice for baseline and n = 6 mice for dicumarol. Student’s t-test, n.s.: not significant.

